# Protein tertiary structure modeling driven by deep learning and contact distance prediction in CASP13

**DOI:** 10.1101/552422

**Authors:** Jie Hou, Tianqi Wu, Renzhi Cao, Jianlin Cheng

## Abstract

Prediction of residue-residue distance relationships (e.g. contacts) has become the key direction to advance protein tertiary structure prediction since 2014 CASP11 experiment, while deep learning has revolutionized the technology for contact and distance distribution prediction since its debut in 2012 CASP10 experiment. During 2018 CASP13 experiment, we enhanced our MULTICOM protein structure prediction system with three major components: contact distance prediction based on deep convolutional neural networks, contact distance-driven template-free (*ab initio*) modeling, and protein model ranking empowered by deep learning and contact prediction, in addition to an update of other components such as template library, sequence database, and alignment tools. Our experiment demonstrates that contact distance prediction and deep learning methods are the key reasons that MULTICOM was ranked 3rd out of all 98 predictors in both template-free and template-based protein structure modeling in CASP13. Deep convolutional neural network can utilize global information in pairwise residue-residue features such as co-evolution scores to substantially improve inter-residue contact distance prediction, which played a decisive role in correctly folding some free modeling and hard template-based modeling targets from scratch. Deep learning also successfully integrated 1D structural features, 2D contact information, and 3D structural quality scores to improve protein model quality assessment, where the contact prediction was demonstrated to consistently enhance ranking of protein models for the first time. The success of MULTICOM system in the CASP13 experiment clearly shows that protein contact distance prediction and model selection driven by powerful deep learning holds the key of solving protein structure prediction problem. However, there are still major challenges in accurately predicting protein contact distance when there are few homologous sequences to generate co-evolutionary signals, folding proteins from noisy contact distances, and ranking models of hard targets.

## 1. Introduction

The major breakthrough in protein structure prediction, particularly template-free (*ab initio*) prediction, is the drastic improvement of the accuracy of residue-residue contact distance prediction in the recent years, leading to the correct folding of some template-free modeling (FM) targets in CASP11 and CASP12 experiment ^1–4^. The accurate prediction of inter-residue contacts and distances has become a key intermediate step and driving force to predict protein three-dimensional (3D) structure from sequence. The breakthrough in contact distance prediction was driven by two key advances: residue-residue co-evolutionary analysis popularized in ^5^ and demonstrated in CASP11 and CASP12 experiment ^4,6^ and deep learning introduced in ^7^ and enhanced in ^8–12^.

The co-evolutionary analysis is based on the observation that two amino acids in contact (or spatially close according to a distance threshold such as 8Å) must co-evolve in order to maintain the contact relationship during evolution, i.e. if one amino acid is mutated to a positively charged residue, the other one must change to a negatively charged one to be in contact. A number of co-evolutionary methods of calculating direct rather than indirect/accidental correlated mutation scores has been developed and shown to improve contact prediction ^13–16^. Moreover, the co-evolutionary scores can be used as input for machine learning methods to further improve contact prediction. Deep learning, the currently most powerful machine learning method, was introduced into the field in 2012 and demonstrated as the best method for protein contact prediction in 2012 CASP10 experiment ^7^. Different variants of deep learning methods - convolutional neural networks and residual networks - were combined with co-evolutionary features to substantially improve contact prediction ^8–12^. The improved contact prediction led to the significant improvement of template-free modeling in CASP12 experiment, in which contact predictions were used with different *ab initio* modeling methods such as fragment assembly and distance geometry to build protein structural models from scratch ^1^.

To prepare for 2018 CASP13 experiment, we focused on enhancing our MULTICOM protein structure prediction system ^17–19^ with our latest development in contact distance prediction empowered by deep learning and its application to template-free modeling and protein model ranking ^17,20–22^, while having a routine update on its other components such as template library, template identification, and template-based modeling. Our experiment demonstrates that contact distance prediction empowered by the advanced deep learning architecture can accurately predict a large number of contacts for some template-free or hard template-based targets, which are sufficient to fold them correctly by the distance geometry and simulated annealing from scratch without using any template or fragment information. Our experiment also shows that directly translating predicted contacts into tertiary structures by satisfying distance restraints can fold large proteins with complicated topologies better than using contacts indirectly to guide traditional fragment assembly approaches. Moreover, we demonstrate that deep learning can integrate 1D, 2D and 3D structural features to improve protein model ranking. Particularly, we show that, for the first time, improved contact prediction can consistently improve protein model ranking. Therefore, contact distance prediction and deep learning are the key driving force that made our MULTICOM predictor rank third in the CASP13 experiment in both template-based and template-free modeling. The success of MULTICOM human and server predictors (MULTICOM_CLUSTER, MULTICOM-CONSTRUCT and MULTICOM-NOVEL) in CASP13 clearly proves that deep learning holds the key for protein contact distance prediction and folding, even though there are still significant challenges in contact/distance prediction for targets with few homologous sequences, translation of noisy or sparse contact distances into 3D models, and selecting a few good protein structural models from a large pool of low-quality ones for a hard target.

## 2. Materials and Method

In this section, we first provide an overview of the MULTICOM server and human prediction system, followed with the detailed description of several key new components that we added into the MULTICOM system in CASP13, such as the protein contact distance prediction empowered by deep learning, *ab initio* protein structure prediction driven by predicted contact distances, and large-scale protein quality assessment enhanced by deep learning and contacts.

### 2.1 An overview of the MULTICOM system

**Figure 1** is an overview of our MULTICOM server and human prediction systems. Once the server received a target protein sequence, MULTICOM searched it against protein sequence databases such as the non-redundant sequence database to collect its homologous sequences to generate multiple sequence alignments, which were used to build sequence profiles such as Position Specific Scoring Matrices (PSSM) ^23^ and Hidden Markov models (HMM) ^24^. The sequence was also used to predict one-dimensional (1D) structural features including secondary structure, solvent accessibility, and disorder regions ^25–26^.

**Figure 1.**
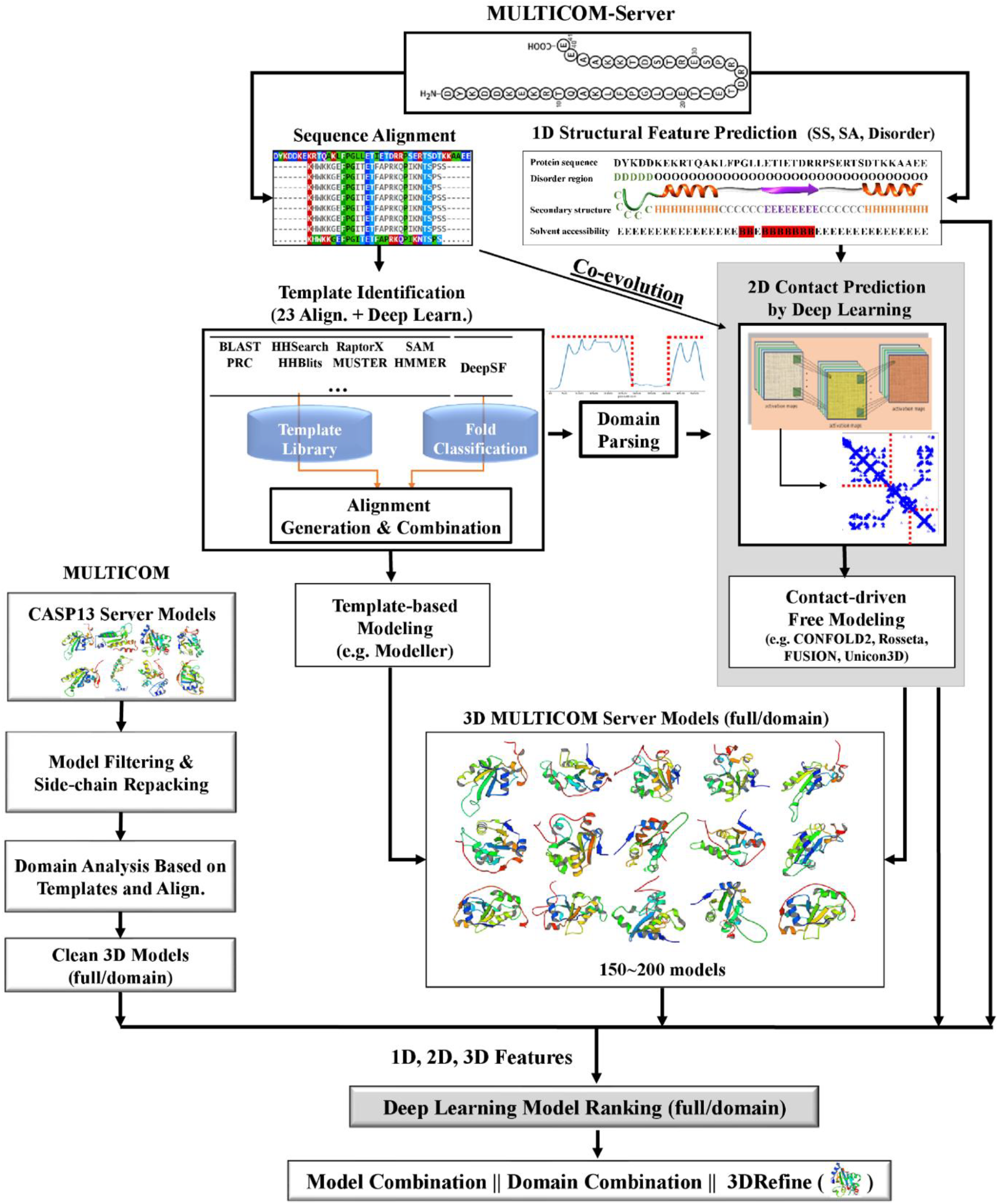
The pipeline of MULTICOM server and human prediction systems.

The profile or sequence of the target was searched against the template profile/sequence library by a number of sequence alignment tools (e.g., BLAST ^27^, CSI-BLAST ^28^, PSI-BLAST ^23^, COMPASS ^29^, FFAS ^30^, HHSearch ^31^, HHblits ^24^, HMMER ^32^, Jackhmmer ^32^, SAM ^33^, PRC ^34^, RaptorX ^35^) to identify protein templates whose structures were known and build pairwise target-template sequence alignments. DeepSF - a deep learning method of classifying protein sequences into folds was also used to identify templates for the target ^36^.

In parallel to the template identification, the multiple sequence alignments of the target were also used to generate co-evolutionary features by CCMpred ^14^, FreeContact ^37^ and PSICOV ^16^, which were used together with other sequential and structural features such as predicted secondary structure and solvent accessibility as input for DNCON2 ^8^ to predict residue-residue contacts at multiple distance thresholds (i.e. 6 Å, 7.5 Å, 8 Å, 8.5 Å and 10 Å).

The target-template sequence alignment was used to identify domain boundaries, i.e. the region of the target not aligned with any significantly homologous template was treated as a template-free modeling domain, otherwise a template-based domain. The contact prediction for template-free domains was made by DNCON2 and combined with the contact prediction of the full-length target.

The pairwise target-template alignments were combined into the multi-template alignments between the target and the multiple templates if the structures of the templates were consistent. The alignments and the structures of templates were fed into Modeller ^38^ to build the structural models for the target. Generally, more than 100 template-based models were constructed for a target.

In parallel to the template-based modeling, predicted contacts were used with several *ab initio* modeling tools such as CONFOLD2 ^39^, Rosetta ^40^, UniCon3D ^41^ and FUSION ^42^ to build structural models for a template-free target or domain. Both the template-based models and/or template-free models were added into a model pool for model ranking.

The MULTICOM human predictor also used all CASP13 server models as input. The incomplete server models or highly similar models (e.g., GDT-TS > 0.95) from the same server group were filtered out. The side chains of the remaining models were repacked by SCWRL ^43^ in order to have the consistent side chain packing before they were evaluated. If the target was identified as multiple-domain protein, the server models were divided into individual domain models.

The structural models from either MULTICOM human predictor or server predictors were compared with 1D structural features (e.g., predicted secondary structure, solvent accessibility) to generate 1D matching scores and with 2D contacts to generate 2D matching scores (i.e., the percentage of predicted contacts existing in a model of the target). The models were also assessed by a number of 3D quality assessment tools to generate 3D quality scores. The 1D, 2D, and 3D quality scores (features) were used by DeepRank-our deep learning-based model quality assessment tool - to predict the accuracy of the models. This quality assessment method was also applied to individual domains if a target had multiple domains. It is worth noting that our three server predictors used different quality assessment methods for model selection. MULTICOM_CLUSTER ranked models primarily based on pairwise similarity scores between models using APOLLO ^44^, while MULTICOM-CONSTRUCT and MULTICOM-NOVEL selected best five models based on our two new *deep learning*-based model ranking methods (DeepRank and DeepRank_avg, described in details in Section 2.4).

The quality assessment scores were used to rank full-length and/or domain-based models and the top ranked models were selected for model combination and refinement. Each top ranked model was combined with other similar models in the ranked list to generate a consensus model. If the consensus model is not substantially different from the initial model (i.e. GDT-TS > 0.88), it was kept as the final model. Otherwise, it was discarded and 3DRefine ^45^ was used to refine the top ranked model to generate a refined final model.

### 2.2 Deep convolutional neural network for contact distance prediction

We used DNCON2 to generate the 2D contact map for an input sequence ^8^. As shown in **Figure 2**, a target sequence was searched against Uniprot20 database (version: 2016_02) by HHblits ^24^ to collect homologous sequences and generate multiple sequence alignments. If there is not a sufficient number of homologous sequences (e.g., < 5L sequences; L sequence length), the target was further searched against Uniref90 database (released by April 2018) by Jackhmmer ^32^ to collect more homologous sequences whose multiple sequence alignments were combined with the results of HHblits search. The multiple sequence alignments were used by CCMPred ^14^, FreeContact ^37^, and PSICOV ^16^ to generate residue-residue co-evolution features. The pairwise co-evolution features together with other pairwise information (e.g. secondary structure, solvent accessibility, and mutual information for each pair of residues) were stored in the L×L input matrices (L: sequence length or domain length).

**Figure 2.**
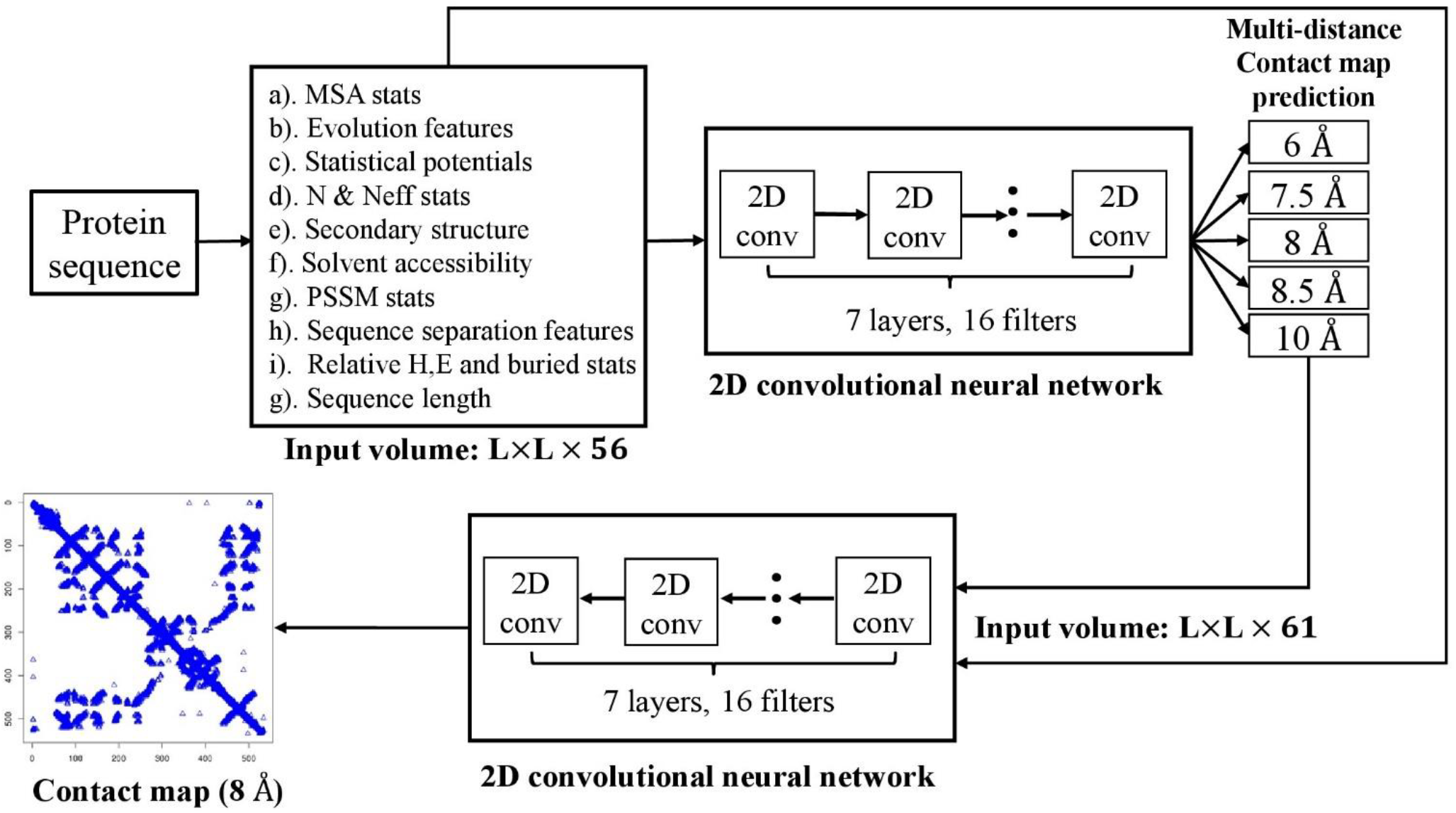
The pipeline of DNCON2 for protein residue-residue contact distance prediction. The input volume has 56 channels (matrices) containing various pairwise residue-residue features.

The input feature matrices were used by the first-level convolutional neural networks in DNCON2 to predict the contact probability maps (i.e. *contact distance distribution*) at multiple distance thresholds 6Å, 7.5 Å, 8 Å, 8.5 Å and 10 Å. The distance distribution and the original input matrices were concatenated as input for the second-level convolutional neural networks to predict a final contact probability map at 8 Å distance threshold.

### 2.3 Contact distance-based *ab initio* folding

We used predicted contacts with a pure contact distance-based *ab initio* modeling tool - CONFOLD2 and several fragment-assembly tools to build 3D models for targets or domains without significant templates being identified. CONFOLD2 ^39^ used only predicted contacts and secondary structures to build structural models without leveraging any other information such as structural fragments **(Figure 3)**. Top *x* × L contacts (*x*: a ratio ranging from 0.1 to 4; L: length of the protein) ranked by probabilities were used to generate distance restraints between *C*_*β*_ atoms (or *C*_*α*_ atom for glycine). The predicted secondary structures were used to generate torsion angle restraints, atom-atom distance restraints, and hydrogen-bond restraints ^46^, which were important for building good local secondary structures in the model. These restraints were used by the distance geometry and simulated annealing optimization implemented in CNS ^47^ to build tertiary structure models by satisfying the restraints as well as possible. In this round of modeling, some local structures, particularly beta-sheets, are often not well formed due to lack of restraints or noisy restraints. To remedy the problem, the potential beta-sheets were detected in the models generated by the first round of modeling. More angular, hydrogen bond, and atom-atom distance restraints were added in order to improve the pairing between the beta strands. Moreover, the contact distance restraints that were not realized in the models were removed from the list. The new set of restraints were used by the distance geometry again to build 3D models. Usually, a few hundred of models were constructed by using different numbers of contact distance restraints (i.e. 0.1L, 0.2L,…., 3.9L, 4L), which were then clustered. Top models from the clusters were selected as final models. The key feature of this approach is that contacts play a dominant and direct role in building structural models. If there are a sufficient amount of accurate distance restraints, high-quality 3D models can be constructed.

**Figure 3.**
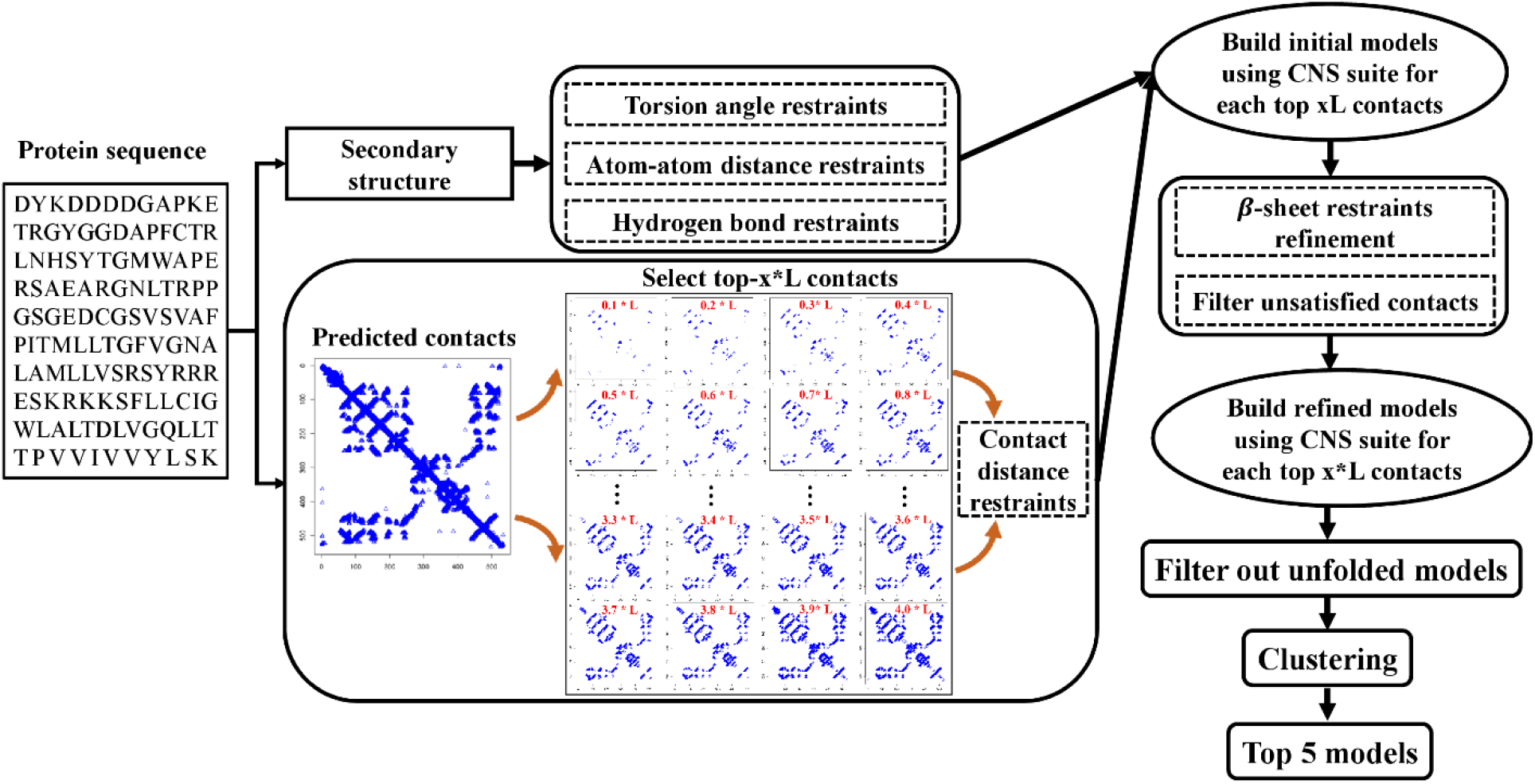
Automated contact distance-based *ab initio* protein structure prediction by CONFOLD2.

As an alternative, we also used predicted contacts as distance or contact restraints with three fragment assembly methods – Rosetta ^40^, UniCon3D ^41^, and FUSION ^42^ to build models. Contacts were used as a part of the energy function of these methods to guide the assembly of protein structure. Rosetta used the structure fragments drawn from a fragment library to assemble the structure, while UniCon3D and FUSION used hidden Markov models to generate conformations for fragments of variable length. In contrast to the CONFOLD approach ^39,46^, extra information such as fragments and energy terms is used in this kind of approach, in which contacts only play an indirect or auxiliary role in structural modeling. Therefore, the fragment assembly approach may fail if its conformation sampling cannot generate correct topologies, which often happens for relatively larger proteins with complicated topologies, even though there is a good amount of accurately predicted contacts. To assist the fragment-assembly with contacts, we selected top L/5 predicted contacts of short-range, medium-range and long-range, which were translated into the distance constraints between pairs of Cβ - Cβ as additional energy terms. Rosetta and FUSION used the bounded potential for a distance *d*, which is defined as follows:

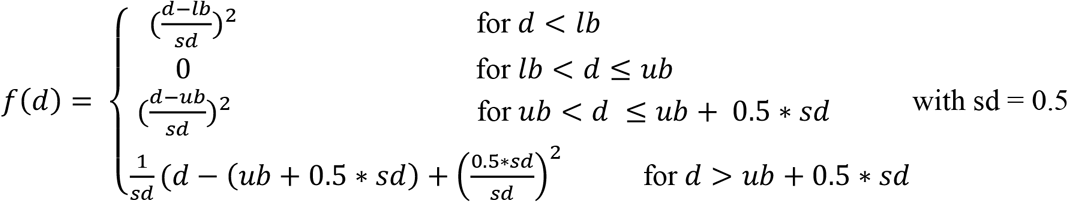

The parameters “*lb*” and “*ub*” are lower and upper bounds for atom-atom distance, which had been optimized and set to 3.5 Å and 8 Å in our experiment. Unicon3D adopted a square well function with the exponential decay to account for the contact distance energy and is defined as:

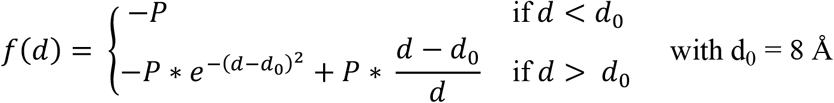

where P is the predicted contact probability for a pair of atoms. In CASP13, the contact-based *ab initio* structure prediction was run for up to two days to generate decoys for model selection.

### 2.4 Protein model ranking by DeepRank integrating 1D, 2D and 3D features

To select most accurate models from a set of predicted structures, we developed a *deep learning*-based quality assessment (QA) method, DeepRank, by integrating multiple QA methods and contact predictions for predicting the global quality of models. Given a pool of models, it first generated one-dimensional (1D) features representing the similarity between the secondary structure and solvent accessibility predicted from the protein sequence by SSPro ^25^ and the ones parsed from each protein model by DSSP ^48^. The percentage of inter-residue contacts (i.e. top L/5 short-range, medium-range and long-range contacts, respectively) predicted by DNCON2 ^8^ existing in a model was used as 2D contact features. It also generated 3D quality scores for each model by using 9 single-model QA methods (i.e. SBROD ^49^, OPUS_PSP ^50^, RF_CB_SRS_OD ^51^, Rwplus ^52^, DeepQA ^22^, ProQ2 ^53^, ProQ3 ^54^, Dope ^55^ and Voronota ^56^) as well as three multi-model QA methods (i.e. APOLLO ^44^, Pcons ^57^, and ModFOLDclust2 ^58^). These features were used by two-level neural networks to predict the quality scores of the models (**Figure 4**). In the first level, all the 1D, 2D and 3D quality features were fed into 10 pre-trained neural networks to predict the quality (GDT-TS score) of each model. These networks were trained on the models of CASP8-11 experiments and rigorously benchmarked on the CASP12 targets. Ten pre-trained neural networks were obtained from 10-fold cross-validations. All the input features of each model were fed into the 10 trained networks to generate 10 quality scores. In the second level, the 10 predicted quality scores and the initial input features were used together by another deep neural network to predict the final quality score. The details of the network configuration are reported in supplemental **Table S5**. This method was also blindly tested as ‘MULTICOM_CLUSTER’ in the CASP13 quality assessment category and ranked as one of the best predictors in selecting models and estimating the absolute error. We also developed a simplified DeepRank method (called DeepRank_avg) by averaging the predictions from the 10 trained networks in the first level as the final quality score.

**Figure 4.**
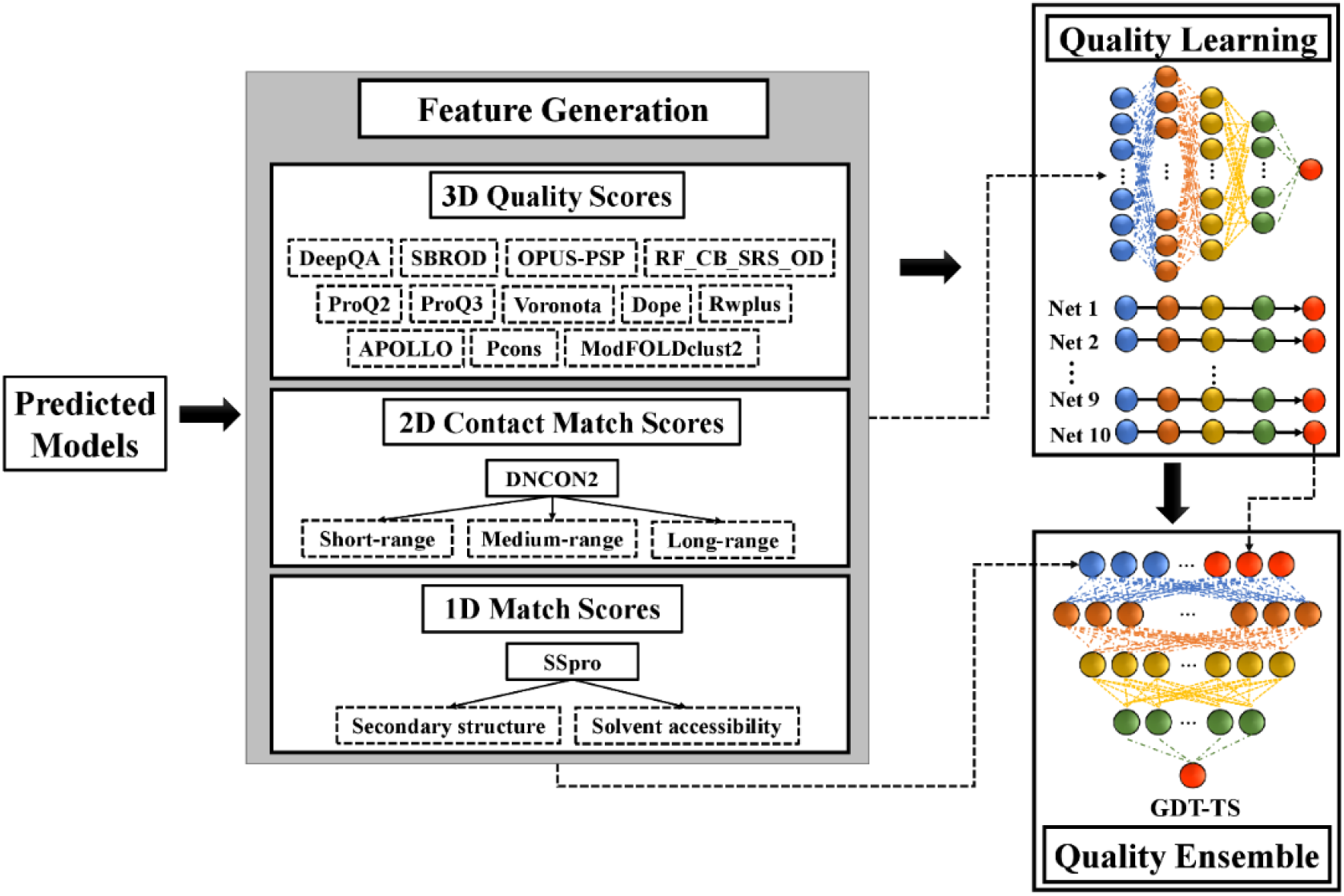
The workflow of *deep learning*-based model quality assessment with contacts (DeepRank).

## 3. Results and Discussions

### 3.1 Performance of MULTICOM human and server predictors in CASP13

We evaluate the performance of MULTICOM methods on 104 “all groups” domains that were used in CASP13 official evaluation. Based on the official domain definition of CASP13, the 104 domains were classified into 31 free-modeling (FM) domains, 40 template-based easy (TBM-easy) domains, 21 template-hard (TBM-hard) domains, and 12 FM-TBM domains.

**Figure 5** shows the performance of MULTICOM human predictor and our three server predictors based on the TM-score metric ^59^. According to the evaluation, as shown in **Figure 5(A)**, MULTICOM human predictor outperforms the three server predictors and MULTICOM-CONSTRUCT ranked better than MULTICOM_CLUSTER, followed with MULTICOM-NOVEL in terms of averaged TM-score on 104 domains. On all the domains, the average TM-score of MULTICOM is 0.69, significantly higher than 0.59 of MULTICOM-CONSTRUCT (difference = 0.1; P-value = 4.478E-14), whereas the difference between the two on template-based easy domain (i.e. 0.04) is much smaller and on template-free domains (i.e. 0.19) is much larger. **Figure 5(B)** shows the performance of four predictors on the 40 TBM-easy domains. MULTICOM-CONSTRUCT and MULTICOM-NOVEL achieved higher TM-score than MULTICOM_CLUSTER. The major difference among the three servers is the QA methods employed for model selection. The three QA methods: DeepRank, DeepRank_avg and APOLLO (a pairwise model comparison method) were used in the MULTICOM_CONSTRUCT, MULTICOM-NOVEL and MULTICOM_CLUSTER, respectively. As shown in supplemental **Figure S5**, DeepRank has the higher capability of model selection than APOLLO. Especially for the template-based targets, DeepRank has a much lower loss (GDT-TS score 0.039) compared to the APOLLO’s loss (0.059) in model selection. The better ability of model selection in template-based targets led to better tertiary structure prediction for MULTICOM-CONSTRUCT (∑ GDT-TS = 75.83) than MULTICOM_CLUSTER (∑ GDT-TS = 72.91) as shown in supplemental **Figure S2**. **Figure 5(C)** reports the results of the four predictors on the 31 free-modeling domains. MULTICOM human predictor successfully predicted correct fold for 17 out of 31 domains (TM-score > 0.5).

**Figure 5.**
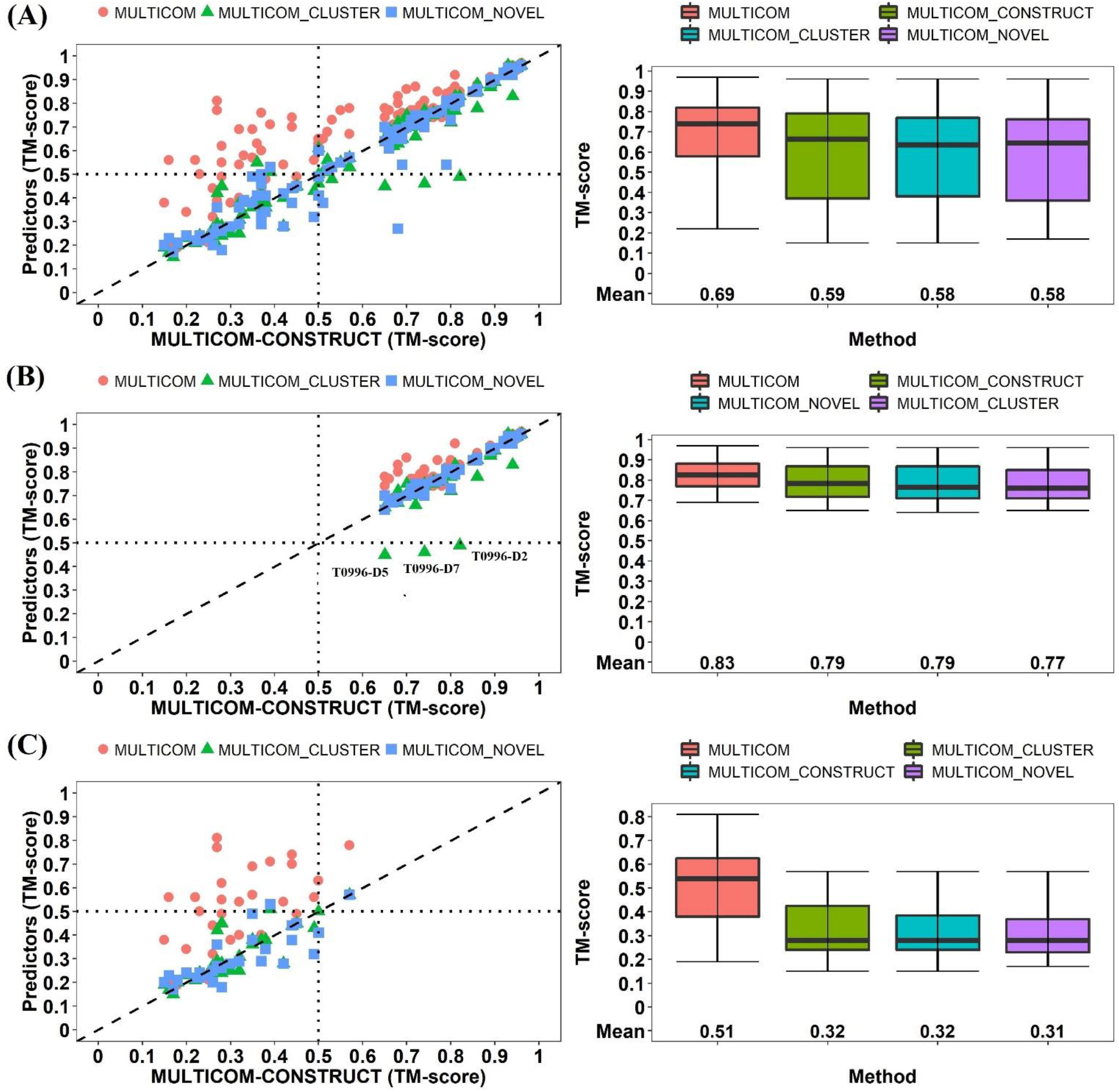
Evaluation of four MULTICOM predictors. The methods are ranked by average TM-score of the first (i.e. TS1) submitted models. **(A)** on 104 domains (Left plot: TM_scores of MULTICOM, MULTICOM_CLUSTER, MULTICOM-NOVEL models versus TM_scores of MULTICOM-CONSTRUCT models; Right plot: mean and variation of the TM-scores of the models of the four methods). **(B)** on 40 template-based (TBM-easy) domains. **(C)** on 31 template-free (FM) domains.

Supplemental **Figure S1** compares MULTICOM with other top ranked CASP13 groups. MULTICOM (group number: ‘089’) is consistently ranked among the top three predictors according to all metrics on the three domain sets. For instance, it is ranked no. 3 according to z-score on all 104 domains. **Figure S2** shows the performance of our three MULTICOM server predictors and other top ranked server groups on the 112 “all groups” and “server only” domains. MULTICOM-CONSTRUCT ranked 7^th^ among all server groups on all the targets, followed by MULTICOM_CLUSTER and MULTICOM-NOVEL. The performance of the global and local quality metrics defined by GDT-TS ^59^, and LDDT score ^60^ are also summarized in **Figure S3** and **Figure S4**.

### 3.2 Performance of DeepRank and individual QA methods used by MULTICOM

To assess how well the model ranking component of MULTICOM predictors worked, we evaluate the results of DeepRank and the individual QA methods used by DeepRank on the CASP13 targets. The loss of each QA method on the 74 CASP13 “all group” full-length targets whose experimental structures are available was calculated and visualized in **Figure 6(A)**. The loss is defined as the difference between the GDT-TS score of the top selected model by each method and the GDT-TS score of the best model of the target. The lower average loss represents the better capability of a QA method for model selection. 24 QA methods are categorized into four groups, including (1) our deep learning integration of diverse quality assessment methods (DeepRank), (2) 3 contact match scores, (3) 3 clustering-based methods, and (4) 17 single-model QA methods. The results show that DeepRank had the lower average loss (0.052) than other individual QA methods on all 74 all-group targets. **Figure 6(B)** plots the GDT-TS scores at the 100-point scale of the top models selected by each individual QA method and DeepRank against the GDT-TS scores of MULTICOM’s first submitted models. The fitted curve for each method is highlighted in different colors. The larger area under the curve represents the better overall accuracy of model selection. The analysis shows that DeepRank achieves higher GDT-TS scores (Avg. GDT = 54.90 at 100-point scale, i.e. 0.549 at 1-point scale) for model selection than the clustering-based method APOLLO (Avg. GDT = 53.31 at 100-point scale, i.e. 0.5331 at 1-point scale), and also outperforms all other QA methods.

**Figure 6.**
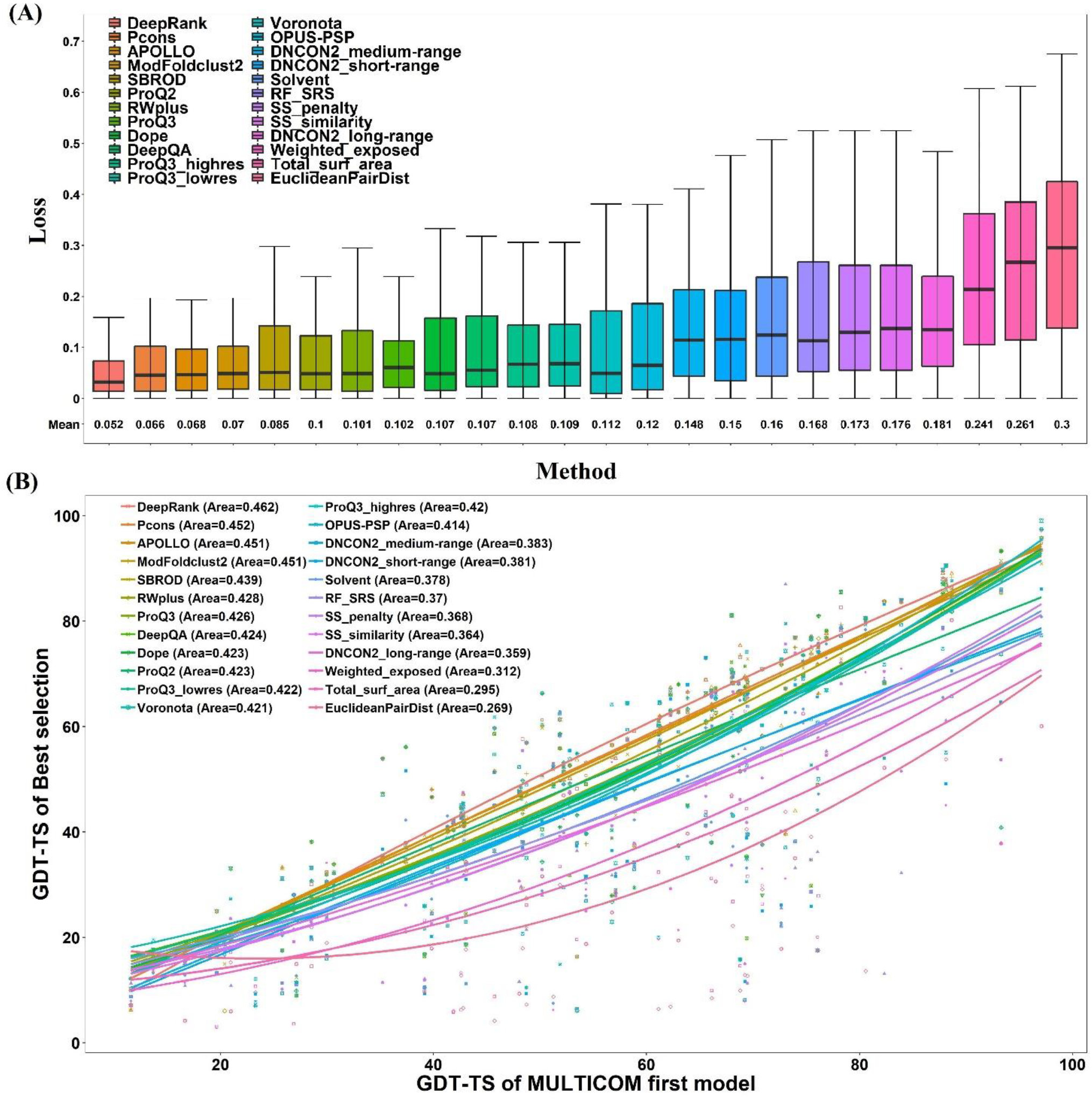
Comparison of DeepRank with individual QA methods used in MULTICOM predictors. (A) The box plot of loss of each method. Here the loss is measure at 1-point scale (i.e. the highest/perfect GDT-TS score = 1). (B) The GDT-TS score at the 100-point scale of the top models selected by each individual QA method and DeepRank is plotted against the GDT-TS score of MULTICOM’s first submitted models for 74 “all group” full-length targets. The curve for each method is fitted by the second-degree polynomial regression function. The area under the curve for each method is calculated and shown on the top left. The larger area indicates the better capacity of model selection.

Prior to CASP13, we assessed how much the deep learning and contact prediction improved the quality assessment in CASP12 dataset. After the quality scores were generated using individual QA methods, two baseline combination strategies (e.g., the average score of raw feature scores and Z-scores respectively) were compared with the deep learning. Supplemental **Table S2** shows that the Z-score based consensus worked better than the average score consensus, while the deep neural network of integrating all features except contacts further reduced the loss from 0.064 of the z-score based consensus to 0.054. Furthermore, the deep learning with contact features performed best (correlation = 0.853 and loss = 0.048), and the improvement was significant compared to the averaging approach (loss = 0.067) according to the P-value (0.007751). The improvement is also consistent with the results in the blind CASP13 experiment (supplemental **Table S3**). The average loss of the deep learning with contacts is 0.051 on the 74 CASP13 targets, lower than 0.059 of the deep learning without contacts that is lower than both the average score consensus and z-score consensus. This further validated the deep learning and contact prediction’s positive contribution to model selection.

**Figure 7** illustrates how MULTICOM estimated the quality of models for a TBM-hard target T0966 and predicted the final structure. **Figure 7(A)** visualized the distribution of the GDT-TS scores of 146 server models for this target. It is a bimodal distribution, where the GDT-TS scores of major models are centered around 0.1 and 0.5. **Figure 7(B)** is the plot of the true GDT-TS scores of models against their predicted ranking by DeepRank. It successfully ranked the model with highest GDT-TS score (0.6103) as No.1 (**Figure 7(D)**). MULTICOM generated a refined model by combining the top 1 selected model with the other top ranked models, which had a GDT-TS score of 0.6113 (**Figure 7(E)**). The ranking of individual QA methods for this target is shown in **Figure S9**. The other three such successful cases for DeepRank are also reported in **Figures S7, S8 and S10**.

**Figure 7.**
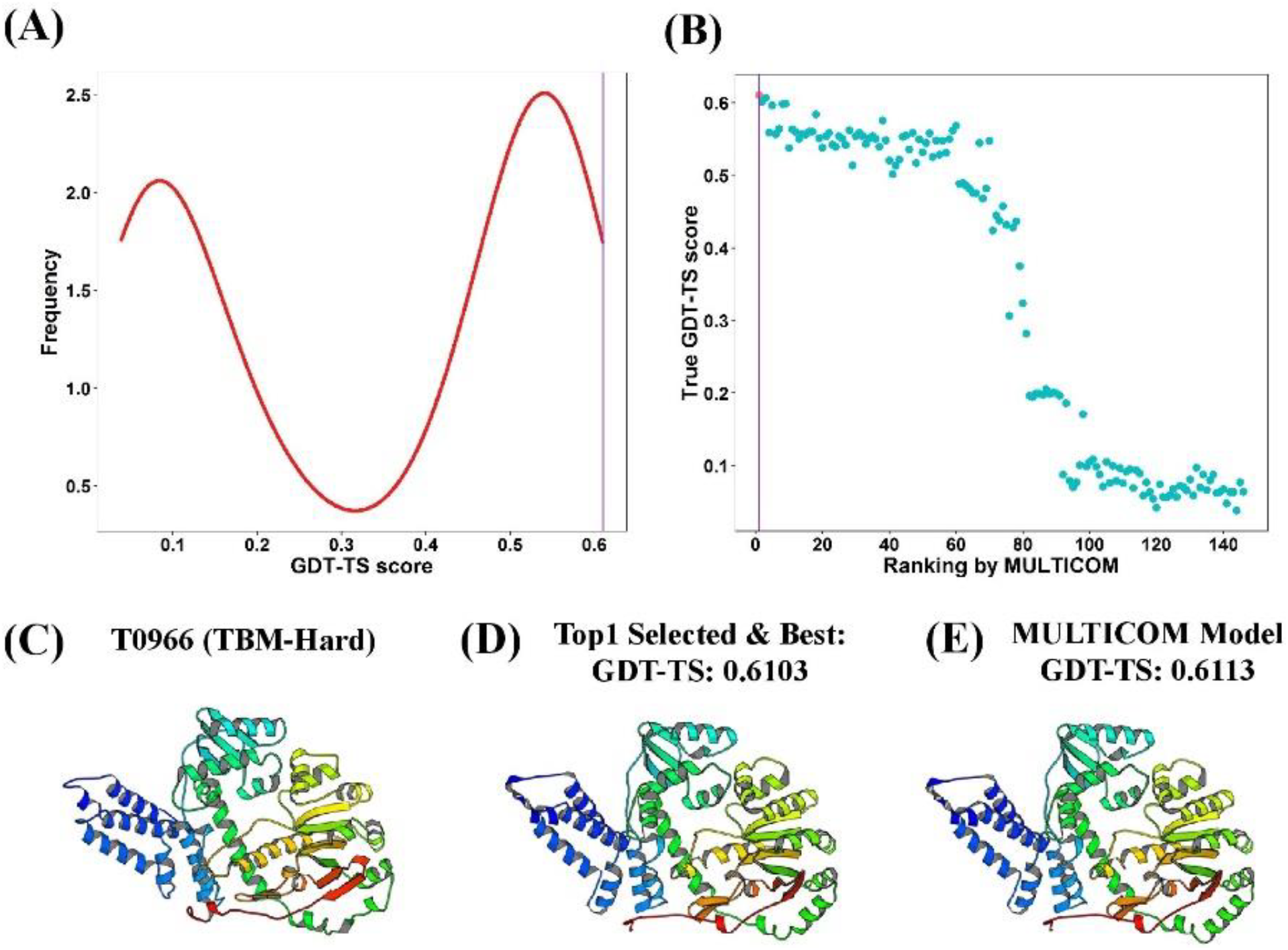
Tertiary structure prediction for T0966. **(A)** The distribution of GDT-TS scores of 146 server models. **(B)** The plot of the true GDT-TS scores of models against their predicted ranking by MULTICOM. The point highlighted in red is the top model selected by DeepRank. **(C)** The native structure of target T0966 (PDB code: 5w6l). **(D)** The top selected model. **(E)** The final first MULTICOM model (TS1).

To assess how contact predictions can help model ranking, we evaluated DeepRank with/without contact features on targets with low contact prediction precision and ones with high contact prediction precision, respectively (**Figure S6**). The consistent, significant improvement in model selection has been observed when the contact prediction of short-range, medium-range, and long-range has high precision (precision > 0.5). However, the less accurate contact prediction led to the slightly worse performance on model selection than not using contact prediction.

### 3.3 Comparison of different contact-based *ab initio* modeling methods on FM targets

To evaluate how predicted contact distances improved template-free modeling, we collected the top 5 models predicted by five *ab initio* modeling methods (CONFOLD2, RosettaCon – Rosetta with contacts, UniCon3D with contacts, FUSION with contacts, and Rosetta without contacts) for all domains that MULTICOM considered them as “hard”. **Figure 8** shows that the GDT-TS scores of the *ab initio* models generally increase as the accuracy of contact prediction becomes higher for each method. This upward trend is most significant for CONFOLD2 and the correlation between the contact accuracy and the GDT-TS score of CONFOLD2 models is 0.578. This is expected because CONFOLD2 is the only pure contact distance-driven modeling method in the group and contact distances play a direct and dominant role in its modeling, while they only play an indirect role in the other three modeling methods assisted by contact predictions.

**Figure 8.**
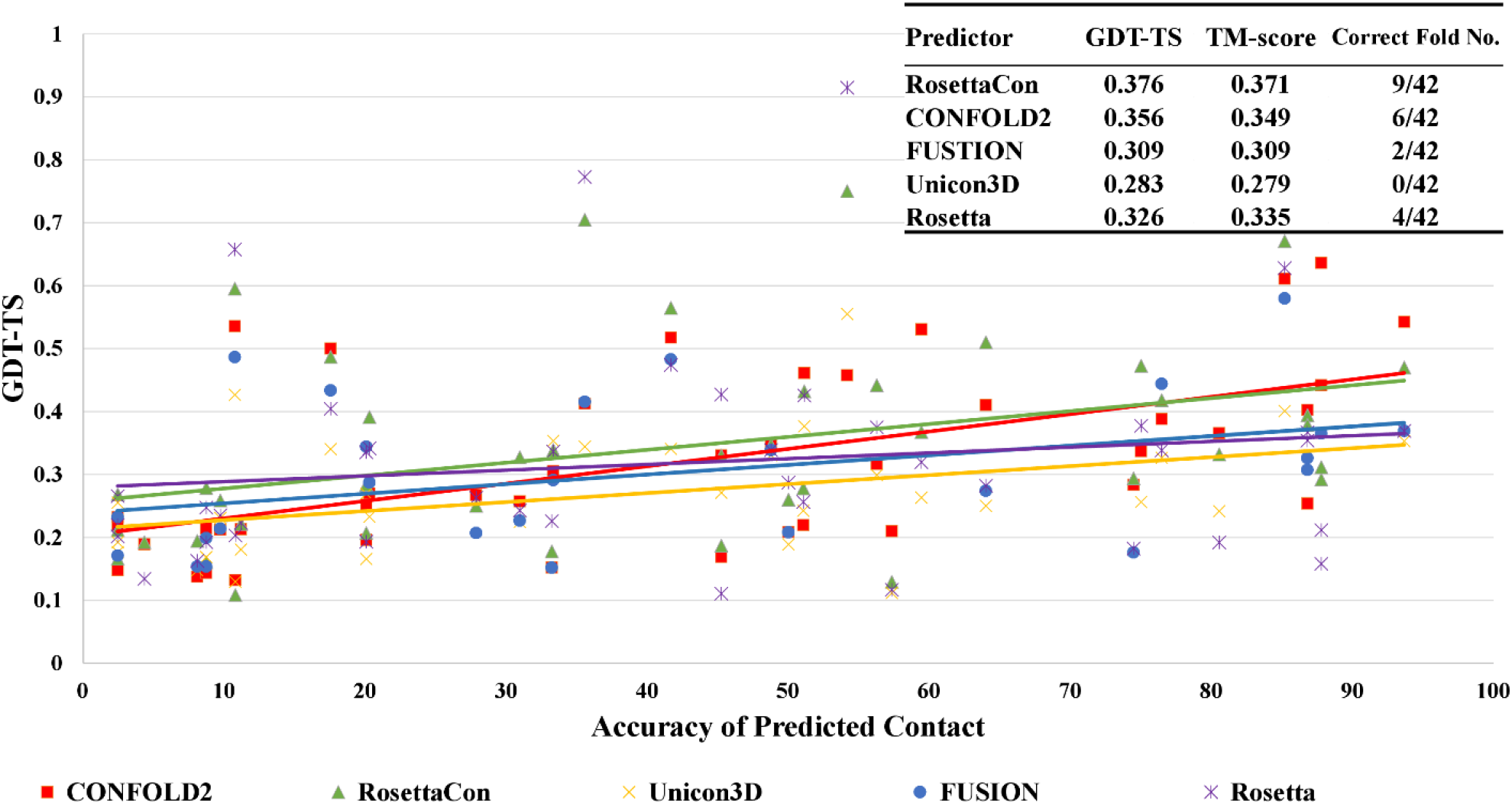
The modeling performance of contact-based *ab initio* modeling methods versus the predicted contact accuracy (L/5 contacts) in CASP13. Each point represents the modeling accuracy in terms of GDT-TS score versus the accuracy of predicted contacts for a method. The colors represent different modeling methods. Rosetta without contacts (purple) was included for comparison. The averaged GDT-TS score and TM-score of five methods on the all CASP13 targets are summarized in the top-right table.

The average GDT-TS score and TM-score were also calculated for each method on the free-modeling targets. The models generated by RosettaCon has the highest average GDT-TS score of 0.376 and CONFOLD2 has the second highest average score of 0.356, followed by Rosetta, FUSION, and UniCon3D. It is interesting to note that CONFOLD2 started to work better than RosettaCon when top L/5 contact predictions reached a high accuracy (e.g. ∼80%). When the accuracy of contact prediction was lower, RosettaCon worked somewhat better than CONFOLD2 probably because the extra structural fragment information and its advanced energy function made some difference. The comparison of RosettaCon and Rosetta shows a 15.3% increase of GDT-TS score by using contact distance restraints, demonstrating that predicted contacts can significantly improve the fragment-assembly modeling.

**Figure 9** show a successful *ab initio* modeling example (a domain of target T1000) for which no significant templates were identified. For the FM domain of T1000 (residues 282-523), the accuracy of top L/5 predicted contacts is 100%, top L 79% and top 2L 50%. CONFOLD2 successfully built a complicated *α*-helix+*β*-sheet+*α*-helix model for the domain with TM-score of 0.8 and GDT-TS of 0.64, while RosettaCon failed to generate a correct topology (i.e. TM-score = 0.33 < 0.5 threshold). This example shows that the pure contact distance driven method such as CONFOLD2 can build high-quality structural models of complicated topology for large domains if a sufficient number of accurate contact predictions are provided.

**Figure 9.**
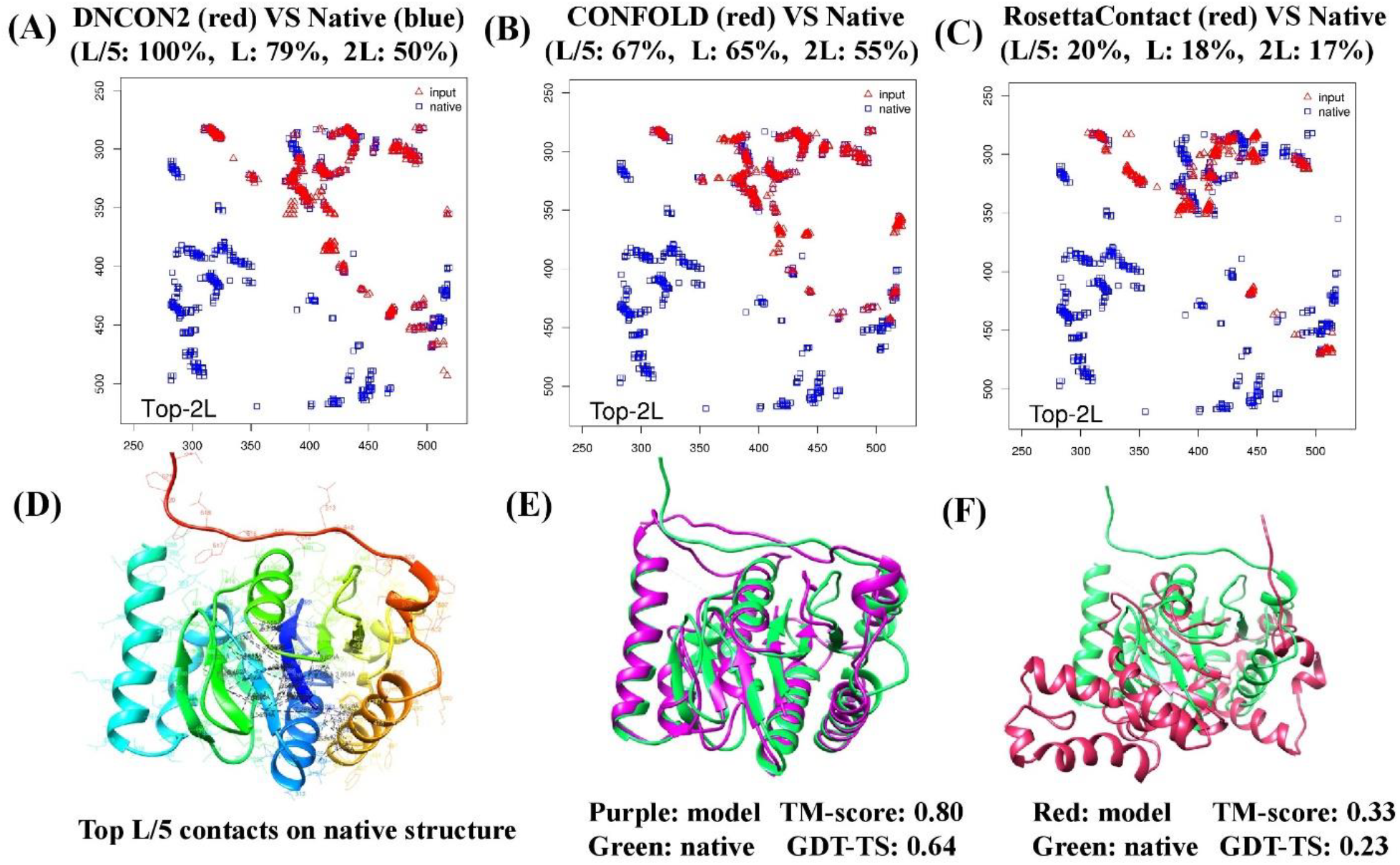
An example of successful contact-driven *ab initio* modeling by CONFOLD2 for a domain of T1000 (residues: 282-523). **(A)** The comparison of the predicted contact map (red, upper triangle) with the true contact map of the native structure (blue). For clear comparison, only the upper triangle of the predicted contact map is shown. The accuracy of predicted contacts is reported at the top of the map. **(B)** The comparison of contact map derived from CONFOLD2 model (red) with the true contact map (blue). The comparison of contact map derived from RossetaCon model (red) with the true contact map (blue). **(D)**The top L/5 contacts visualized in the native structure. **(E)** The superposition of CONFOLD2 model (purple) and the native structure (green). The TM-score and GDT-TS score of the model is shown under the model. **(F)** The superposition of RosettaCon model (red) and the native structure (green).

### 3.4 Impact of domain parsing on structure prediction and model ranking

Protein domain identification is an important component in the MULTICOM predictors. When a target protein sequence was searched against a template library, the domain regions that were homologous to templates were marked as “template-based” and modeled by the template-based modeling protocol. The unmarked regions were modeled by the contact distance-based *ab initio* modeling methods. The domain models were evaluated using the three QA methods and top models were assembled into full-length structures as final predictions. For the human predictor, the domain boundaries might be re-analyzed by taking the structural information of top ranked server models into account. We assessed the impact of domain parsing on the structure prediction of the CASP13 targets that were predicted as multi-domain proteins. The final predicted models of these multi-domain targets and the models without domain parsing were evaluated and compared according to the official domain definitions of CASP13. Among the 90 CASP13 targets, 31 targets were modeled as multi-domain by MULTICOM server predictors and 19 targets by MULTICOM human predictor. Supplemental **Table S6** reports the scores of the models using or not using domain parsing. For the server predictors, the performance of structure prediction was substantially improved in terms of GDT-TS, TM-score and RMSD after the domain-based modeling was applied. For the human predictor, the quality of final predictions was also slightly improved when domain information was considered. And almost all the improvement is significant.

### 3.5 What went right?

In CASP13, a main progress was to apply contact distance prediction and deep learning to improve *ab initio* modeling. Predicted contacts were successfully utilized to guide *ab initio* structure modeling for several hard targets that could never be modeled correctly before. Supplemental **Figure S11** shows the models and scores of nine hard targets that were folded into correct topology when the predicted contacts generated by DNCON2 were rather accurate. Remarkably, a pure contact distance-driven modeling method – CONFOLD2 can correctly predict complex folds of large domains if a sufficient amount of accurate contact distance predictions is provided. Furthermore, the inter-residue distance distribution predicted by DNCON2 (e.g. 6 Å, 7.5 Å, 8 Å, 8.5 Å and 10 Å) is valuable for structure prediction, demonstrated by the fact that it helped improve the accuracy of final top L/5 contact predictions from 57.11% to 61.97% on CASP13 targets (supplemental **Figure S12**).

Another main progress is that MULTICOM performed better in ranking the models in CASP13 than in CASP12 due to the application of deep learning and contact prediction. MULTICOM successfully selected models that are identical or close to the best models for 28 targets (see the distribution of loss of model selection for all the targets and two good examples in supplemental **Figure S13**).

Moreover, we successfully tested a new heuristic method to apply domain-based contact predictions to validate multi-domain template-based models. One such example is T0996, a challenging template-based modeling target due to its very large size and very weak homology with existing templates (**Figure 10**). It was recognized by CASP13 as hard template-based target because only several weak partial templates (e.g. PDB code: 5UW2, chain A) could be detected. MULTICOM server predictors successfully divided T0996 into 7 domains and the predicted domain boundaries were largely accurate compared to the official domain definition. Each domain region was modeled through MULTICOM domain-based modeling pipeline. After the domain models were assembled, the full-length structural model was evaluated by the predicted contacts using ConEva ^61^. The contacts in the model matched well with the contacts predicted by DNCON2 domain by domain, confirming that both domain parsing and structure modeling was largely correct (**Figure 10**). This contact-based validation approach was applied to all CASP13 targets during CASP13, providing a complementary validation for structure modeling.

**Figure 10.**
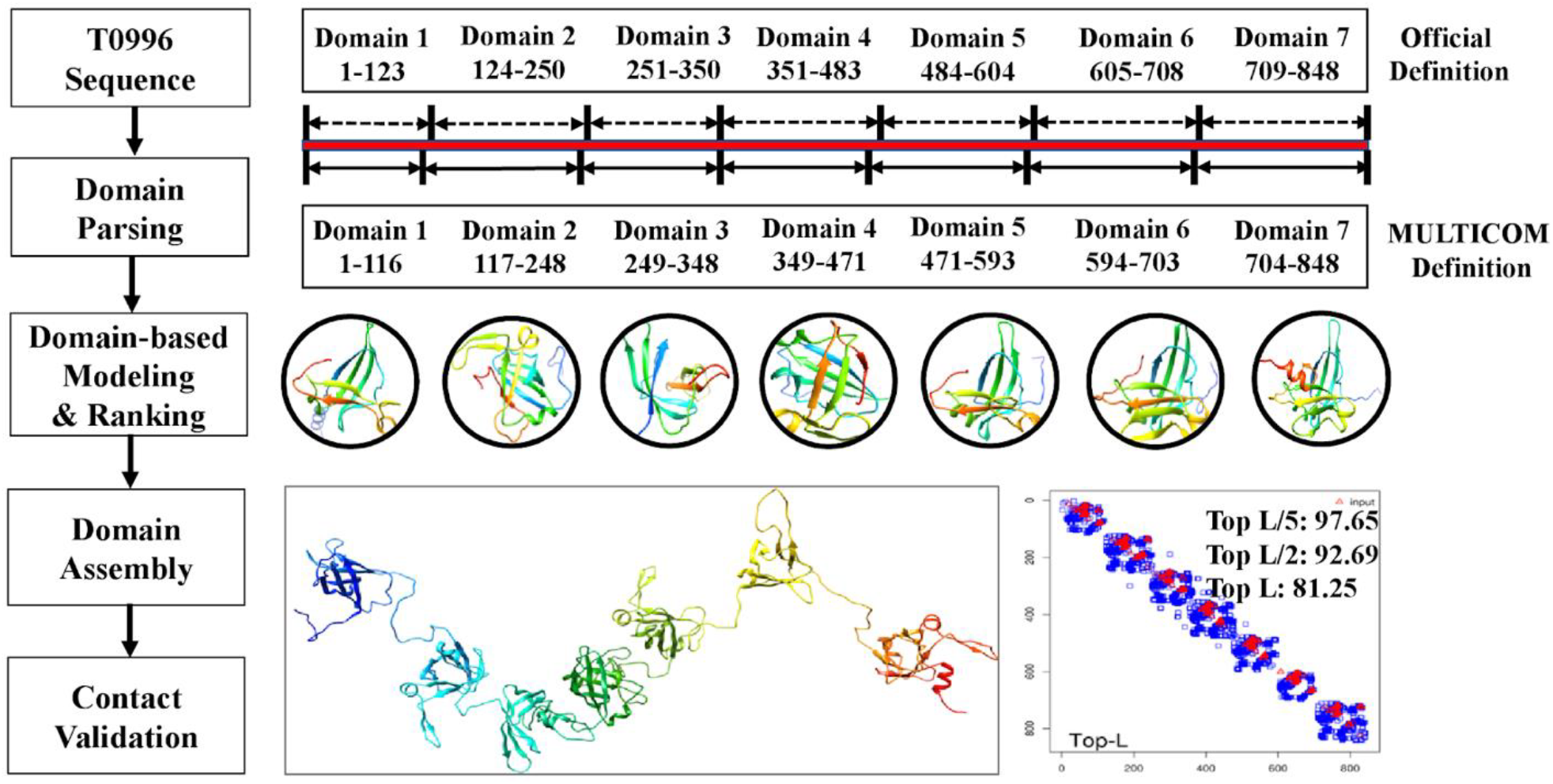
The successful modeling of a large multi-domain target T0996 and the contact-based validation. The contacts (red) predicted by DNCON2 match with the contacts (blue) in the template-based models domain by domain.

### 3.6 What went wrong?

Despite the significant progress of MULTICOM in CASP13, it has its several limitations. The first limitation is in contact distance prediction. DNCON2 sometime failed to generate a sufficient amount of accurate contact predictions to predict correct folds. The problem is particularly severe when the number of effective homologous sequences for a target is small (see supplemental **Figure S14** for an example – T0998). One possible reason is that it did not use a metagenomics sequence database ^62^ that contains sequences not present in the non-redundant protein sequence database and the latest HHblits database ^24^ to collect homologous sequences. Another possible reason is the convolutional architecture used by DNCON2 is not deep enough in comparison with some other approaches ^10,12,63^. The second limitation is that only the coarse distance restraints derived from binary contacts at 8 Å threshold were used with CONFOLD2 for *ab initio* modeling, without taking advantage of the more detailed distance distribution spanning multiple distance thresholds predicted by DNCON2, which limited its capability to build quality models ^64^.

The third limitation is that the deep learning-based quality assessment failed on some targets. As shown in supplemental **Figure S13 (B)**, DeepRank method performed poorly with loss > 0.1 on 14 “all groups” targets. The failed rankings are summarized in supplemental **Table S4** and **Figure S15-S28**. The results show that its performance was worse on the free-modeling targets or hard-template targets than on other targets. A possible reason is that a large portion of low-quality models in the pool and less accurate features of measuring model quality (e.g. contact predictions) for the hard targets hinders the performance of the deep learning ranking.

## 4. Conclusion and Future Work

Our CASP13 results demonstrate that residue-residue contact prediction, more generally distance prediction, is the key direction to advance protein structure prediction, particularly *ab initio* prediction, and deep learning is the key technology to solve it. Not only do accurate contact distance prediction and deep learning enhance *ab initio* structure folding, but also model ranking for both template-based and free modeling. In the future, we will develop more advanced deep learning methods to directly predict real-value distances between residues and/or classify them into much finer intervals than DNCON2 currently does. The more detailed distance predictions will be used to more accurately fold proteins by the distance geometry ^39,46^, simulated annealing and advanced gradient descent optimization ^65–66^ as well as to rank protein models.

## 5. Acknowledgements

This work is supported by an NIH grant (R01GM093123), an NSF IIS grant (IIS1763246), and an NSF DBI grant (DBI1759934) to JC.

