## Supplementary File for "Protein tertiary structure modeling driven by deep learning and contact distance prediction in CASP13"

### Tables

| Rank | GR Code | GR name | Count | Avg. GDT-TS | Sum Zscore (>0.0) | Avg. RMSD | Avg. LDDT | Avg. TM-score |
|---|---|---|---|---|---|---|---|---|
| 1 | 145 | QUARK | 112 | 62.37 | 98.83 | 3.1 | 0.56 | 0.68 |
| 2 | 261 | Zhang-Server | 112 | 62.48 | 97.17 | 3 | 0.57 | 0.68 |
| 3 | 324 | RaptorX-DeepModeller | 112 | 61.19 | 87.57 | 3.41 | 0.56 | 0.67 |
| 4 | 221 | RaptorX-TBM | 112 | 58.66 | 65.99 | 3.35 | 0.54 | 0.64 |
| 5 | 368 | BAKER-ROSETTASERVER | 111 | 55.24 | 51.45 | 2.99 | 0.52 | 0.61 |
| 6 | 498 | RaptorX-Contact | 112 | 53.26 | 46.33 | 4.32 | 0.5 | 0.6 |
| **7** | **243** | **MULTICOM-CONSTRUCT** | **112** | **54.6** | **37.2** | **3.04** | **0.5** | **0.6** |
| **8** | **058** | **MULTICOM_CLUSTER** | **112** | **53.81** | **34.8** | **3.29** | **0.49** | **0.59** |
| **9** | **023** | **MULTICOM-NOVEL** | **112** | **53.79** | **32.26** | **3.5** | **0.5** | **0.59** |
| 10 | 164 | Yang-Server | 109 | 53.4 | 28.78 | 3.02 | 0.46 | 0.59 |

**Table S1.** The official CASP13 evaluation of top 10 server predictors on all 112 “all groups + server only” domains. Predictors were ranked by the sum of Z-score of the first (i.e. TS1) submitted models predicted by all predictors. Z-scores were calculated according to the GDT-TS scores of predicted models. The results of three MULTICOM server predictors (MULTICOM_CLUSTER,

MULTICOM-CONSTRUCT and MULTICOM-NOVEL) are highlighted in the table. All the results are compiled from the official data released at the CASP13 website. These 10 predictors came from five research labs: Zhang Lab (QUARK and Zhang-Server), Xu Lab (RaptorX-DeepModeller, RaptorX-TBM, RaptorX-Contact), Baker Lab (ROSETTASERVER), Cheng Lab (MULTICOM-CONSTRUCT, MULTICOM_CLUSTER, MULTICOM-NOVEL) and Yang Lab (Yang-Server).

| Data | Contact | Correlation | Loss |
|---|---|---|---|
| CASP12 | DeepRank-NoContact | 0.832 | 0.054 |
| | **DeepRank-WithContact** | **0.853** | **0.048** |
| | Average features | 0.764 | 0.067 |
| | Average feature_zscore | 0.720 | 0.064 |
| CASP13 (74 targets) | DeepRank-NoContact | 0.869 | 0.059 |
| | **DeepRank-WithContact** | **0.898** | **0.051** |
| | Average_feature | 0.777 | 0.088 |
| | Average_feature_zscore | 0.757 | 0.068 |

**Table S2.** Deep learning and contact prediction improved protein model quality assessment in CASP12 and CASP 13 datasets.

| | **Loss** | | | **Correlation** | | |
|---|---|---|---|---|---|---|
| **Method** | **All** | **TBM + TBM-hard** | **FM + FM/TBM** | **All** | **TBM + TBM-hard** | **FM + FM/TBM** |
| **DeepRank No-Contact** | 0.059 | 0.046 | 0.066 | 0.869 | 0.936 | 0.824 |
| **DeepRank with Contacts** | **0.051** | **0.040** | **0.061** | **0.898** | **0.963** | **0.857** |

**Table S3.** Impact of contact features on protein model quality assessment in CASP13 dataset.

| **Target** | Classification | Loss | **Target** | Classification | Loss | **Target** | Classification | Loss |
|---|---|---|---|---|---|---|---|---|
| **T0953s2** | FM/TBM+FM | 0.14 | **T0980s2** | FM | 0.11 | **T1019s1** | FM/TBM | 0.14 |
| **T0957s1** | FM+TBM-hard | 0.12 | **T0991** | FM | 0.12 | **T1022s2** | TBM-hard | 0.1 |
| **T0968s1** | FM | 0.16 | **T0992** | FM/TBM | 0.13 | | | |
| **T0975** | FM | 0.2 | **T0998** | FM | 0.16 | | | |
| **T0978** | FM/TBM | 0.14 | **T1008** | FM/TBM | 0.2 | | | |
| **T0979** | TBM-hard | 0.12 | **T1010** | FM | 0.3 | | | |

**Table S4.** CASP13 targets with model selection loss > 0.1 by DeepRank quality assessment (QA) method.

| Stage | Network | Optimization | Activation | Nodes in Hidden Layer |
|---|---|---|---|---|
| Stage 1 | Network 1 | nadam | Sigmoid | 5 |
| | Network 2 | nadam | Sigmoid | 15-15 |
| | Network 3 | nadam | Sigmoid | 5 |
| | Network 4 | nadam | Sigmoid | 70-25 |
| | Network 5 | nadam | Sigmoid | 5-5 |
| | Network 6 | nadam | Sigmoid | 25-15 |
| | Network 7 | nadam | Sigmoid | 85-15 |
| | Network 8 | nadam | Sigmoid | 100-70 |
| | Network 9 | nadam | Sigmoid | 20-5 |
| | Network 10 | nadam | Sigmoid | 20-15 |
| Stage 2 | Network 1 | nadam | Sigmoid | 5 |

**Table S5.** The configuration of Stage 1 and Stage 2 neural networks of DeepRank.

| Predictor | # | GDT-TS | | | TM-score | | | RMSD | | |
|---|---|---|---|---|---|---|---|---|---|---|
| | | Before | After | P-value | Before | After | P-value | Before | After | P-value |
| MULTICOM CLUSTER | 62 | 0.45 | **0.56** | 8.8E-06 | 0.50 | **0.62** | 3.6E-06 | 16.08 | **8.99** | 2.4E-05 |
| MULTICOM CONSTRUCT | 62 | 0.45 | **0.55** | 9.5E-06 | 0.50 | **0.60** | 7.6E-06 | 30.38 | **8.78** | 4.7E-06 |
| MULTICOM (HUMAN) | 48 | 0.64 | **0.66** | 7.8E-03 | 0.67 | **0.69** | 2.3E-02 | 7.75 | **5.93** | 7.7E-02 |

**Table S6**. Effect of domain parsing on the protein structure prediction. The protein targets which were treated as multi-domain proteins in CASP13 by MULTICOM predictors are evaluated. MULTICOM server predictor predicted 31 out of 90 targets as multi-domain proteins, while MULTICOM human predictor modeled 19 targets as multi-domain ones. The second column lists the total number of individual domains of the multi-domain targets. The results show that the quality after domain parsing is always better than not using domain parsing for the three predictors.

### Figures

#### Section I. Performance of MULTICOM structure prediction methods

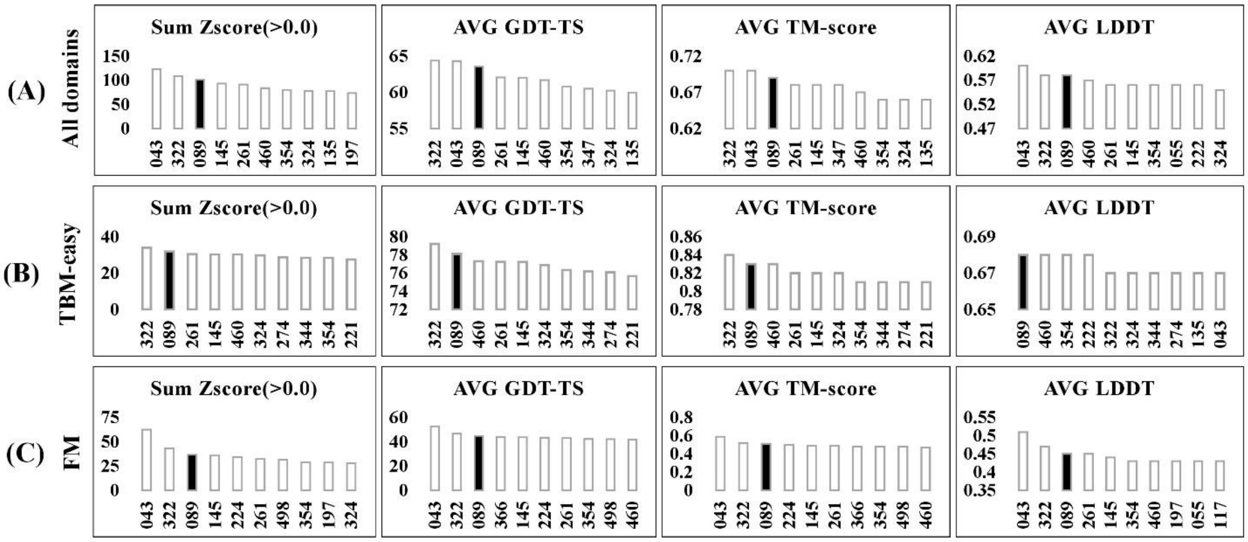


**Figure S1.** The official CASP13 evaluation of top 10 predictors out of 98 human and server predictors on 104 "all groups" targets. Predictors are ranked based on four different metrics of the first (i.e. TS1) submitted models predicted by all predictors. All the results are compiled from the official data released at the CASP13 website. (A) Evaluation on 104 domains. (B) Evaluation on 40 template-based (TBM-easy) domains. (C) Evaluation on 31 template-free (FM) domains. The bar highlighted in black is the performance of MULTICOM human predictor (group number: '089') in CASP13. The figures were drawn according to the results compiled from the official data released at the CASP13 website.

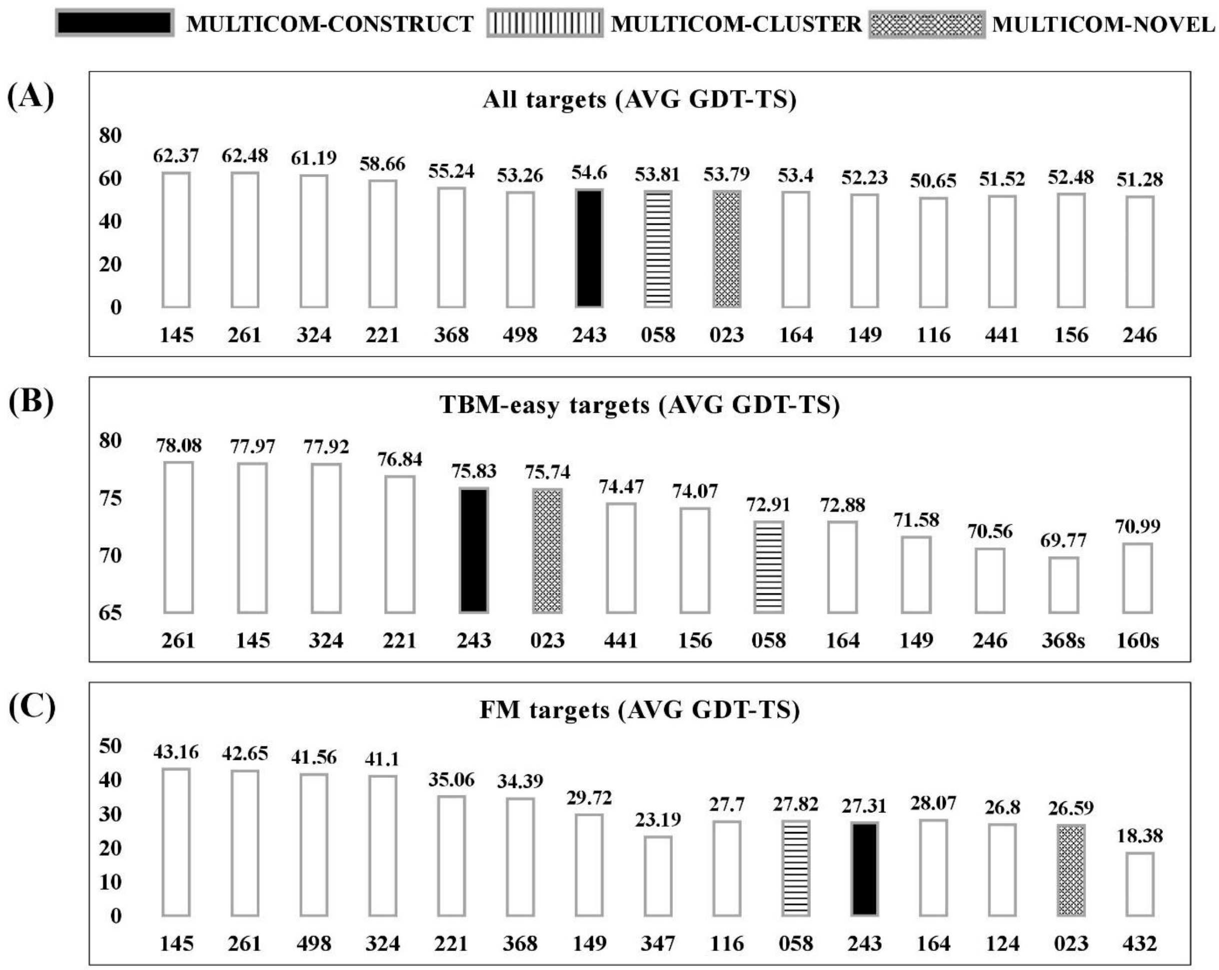


**Figure S2.** The official CASP13 evaluation of top 15 server predictors on 112 “all groups + server only” domains. Predictors are ranked by sum of Z-score of the first (i.e. TS1) submitted models predicted by all predictors denoted by each bar. Z-score (not shown) of each predictor was calculated according to GDT-TS scores of predicted models. The average GDT-TS score at the 100-point scale of each predictor is shown at the top of each bar and the server group code at the bottom. All the results are compiled from the official data released at the CASP13 website. (A) Evaluation on 112 domains. (B) Evaluation on 45 template-based (TBM-easy) domains. (C) Evaluation on 32 template-free (FM) domains. The highlighted bars are the performance of MULTICOM-CONSTRUCT (black filled), MULTICOM-CLUSTER (Horizontal stripes), MULTICOM-NOVEL (Dotted) in CASP13. The detailed results are summarized in **Table S1**.

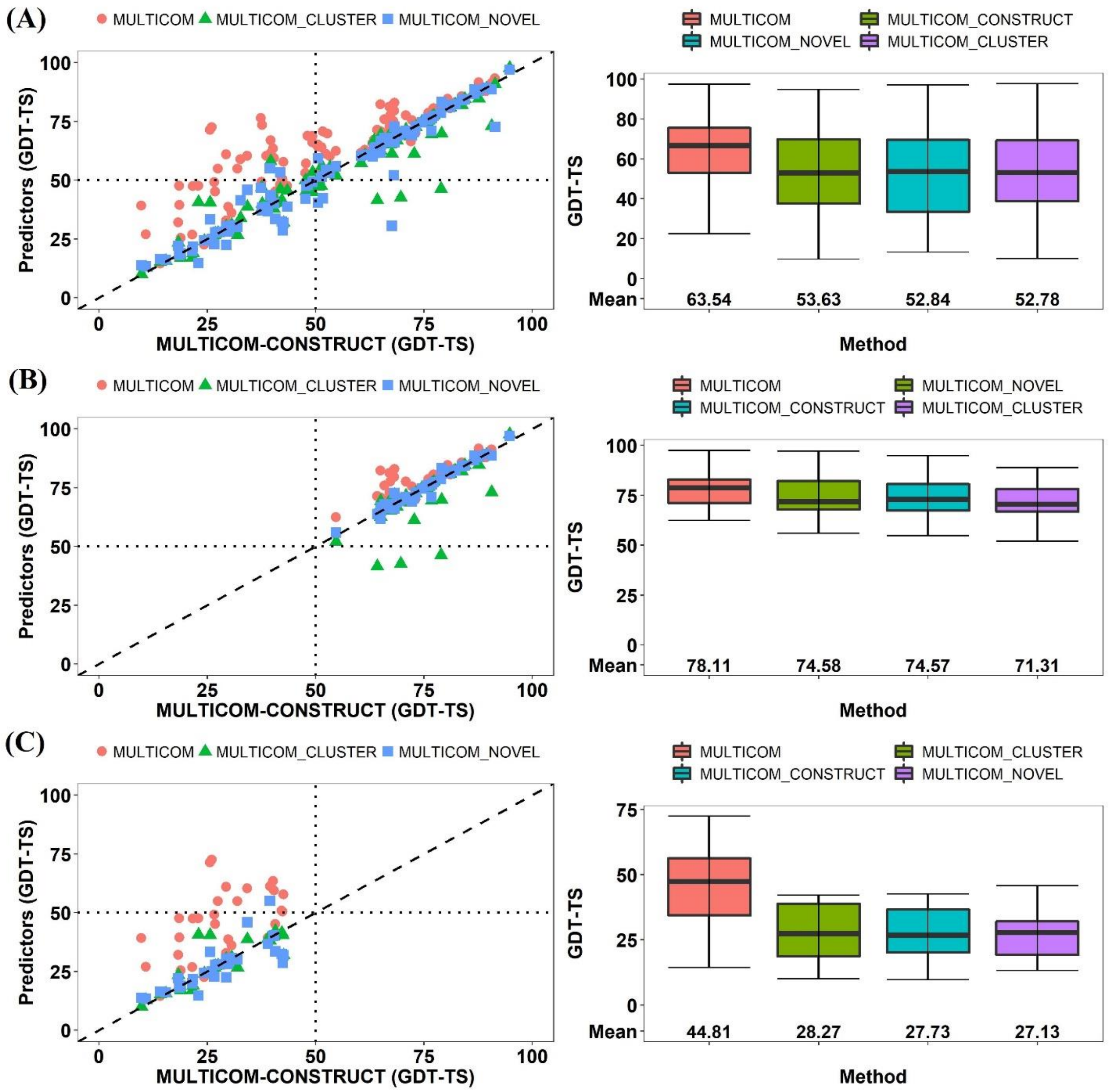


**Figure S3.** Evaluation of four MULTICOM predictors by GDT-TS score. The methods are ranked by average GDT-TS score of the first (i.e. TS1) submitted models. **(A)** on 104 domains (Left plot: GDT-TS scores of MULTICOM, MULTICOM_CLUSTER, MULTICOM-NOVEL models versus GDT-TS scores of MULTICOM-CONSTRUCT models; Right plot: mean and variation of the GDT-TS scores of the models of the four methods). **(B)** on 40 template-based (TBM-easy) domains. **(C)** on 31 template-free (FM) domains.

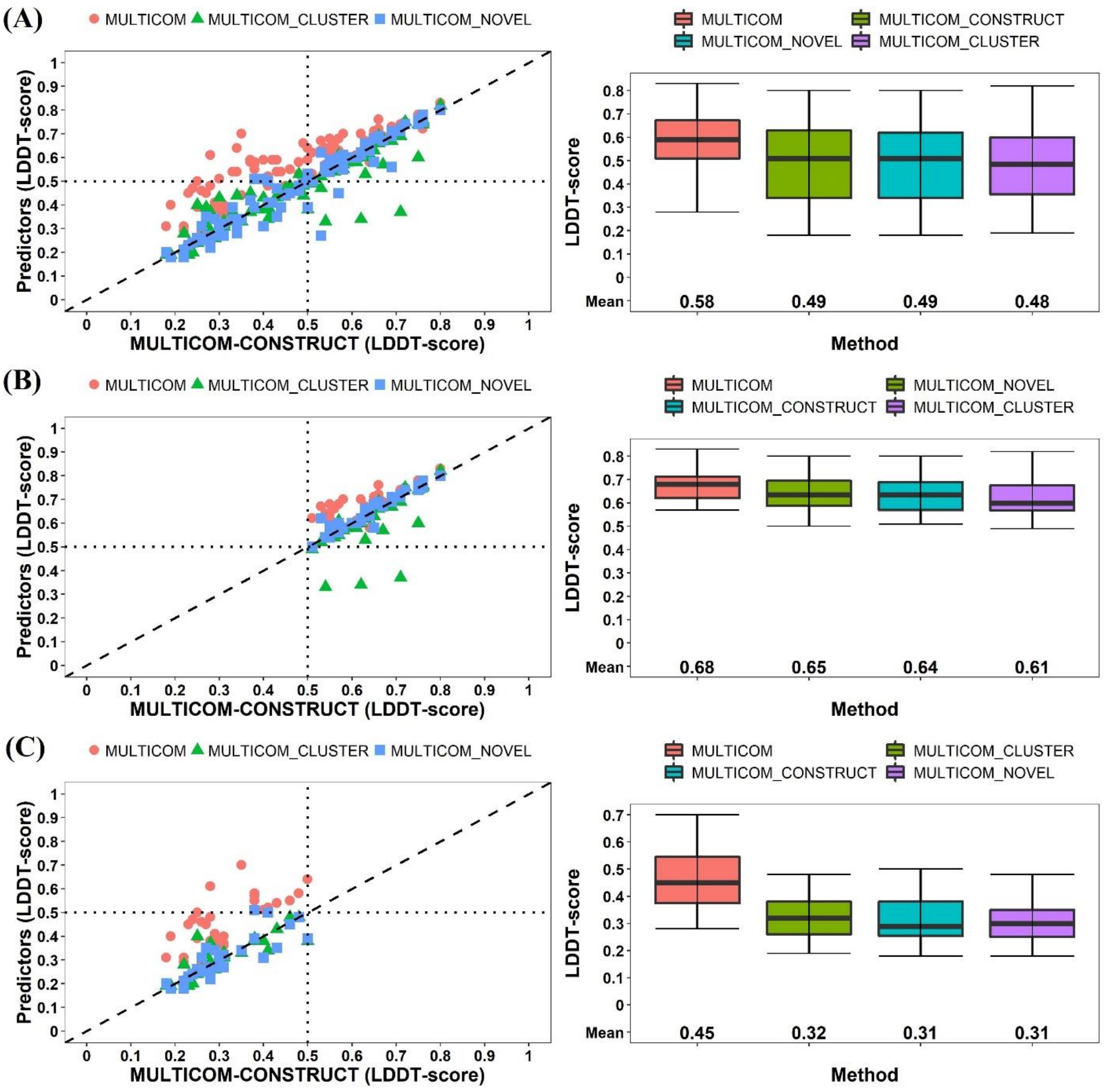


**Figure S4.** Evaluation of four MULTICOM predictors by LDDT score. The methods are ranked by average LDDT score of the first (i.e. TS1) submitted models. **(A)** on 104 domains (Left plot: LDDT scores of MULTICOM, MULTICOM_CLUSTER, MULTICOM-NOVEL models versus LDDT scores of MULTICOM-CONSTRUCT models; Right plot: mean and variation of the LDDT scores of the models of the four methods). **(B)** on 40 template-based (TBM-easy) domains. **(C)** on 31 template-free (FM) domains.

**Section II. Performance of DeepRank method for model selection**

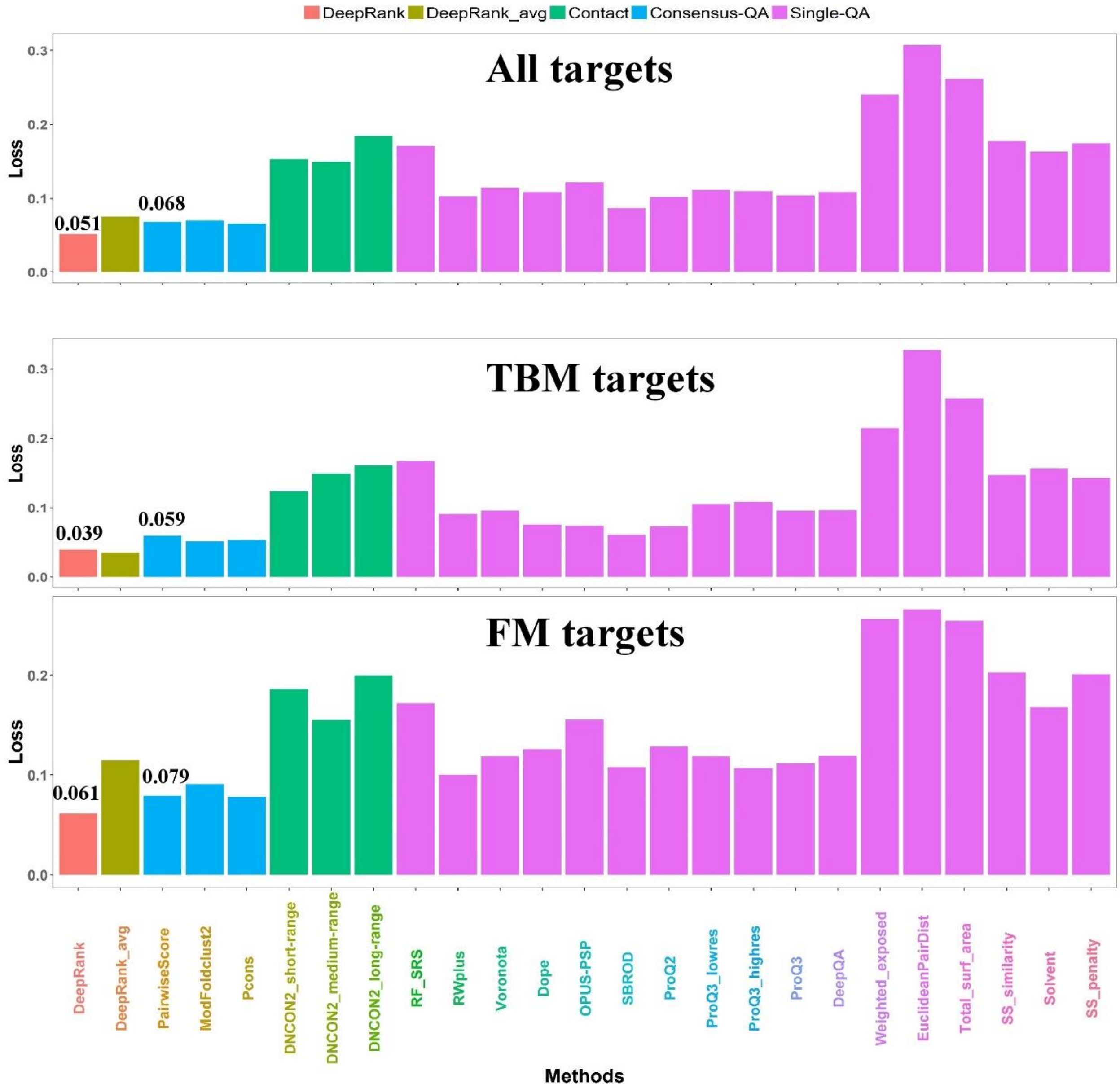


**Figure S5.** Comparison of DeepRank with individual QA features on CASP13 targets. The methods are evaluated according to average GDT-TS loss calculated from the 74 full-length targets, 40 templated-based (TBM-easy and TBM-hard) targets, and 37 free-modeling targets (FM+FM/TBM), respectively.

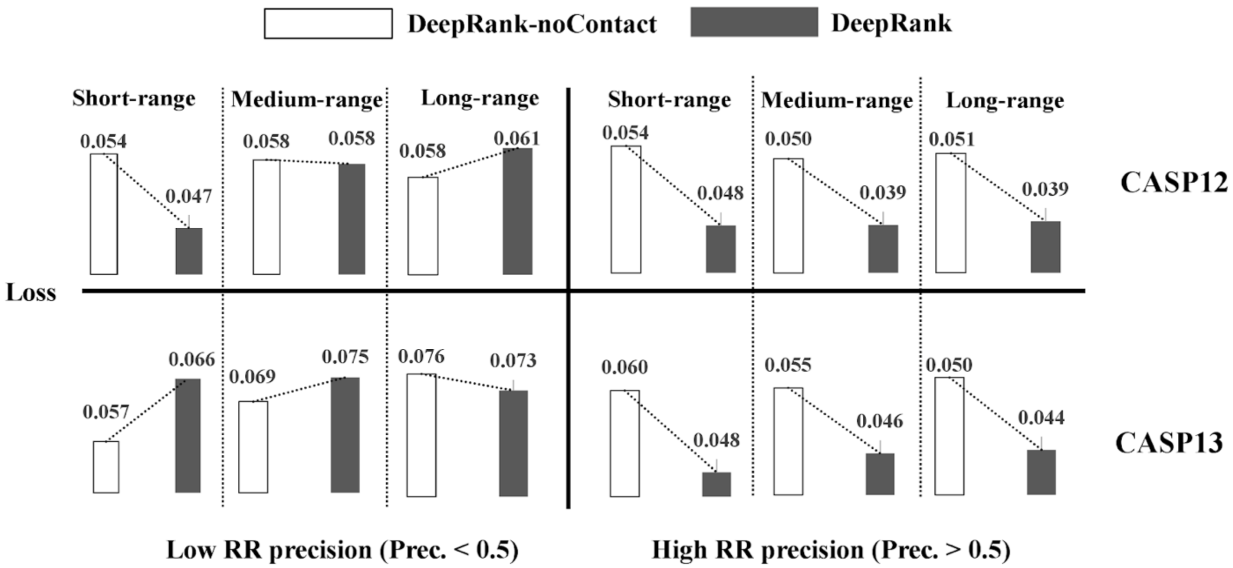


**Figure S6**. Impact of contact prediction accuracy on protein model quality assessment in CASP12 and CASP13 datasets. The white bar represents the loss of DeepRank QA method without contact information, while the black bar represents that of the DeepRank method with contact information. The loss with/without each kind of contact features (short-range, medium-range, long-range) is shown. The loss was consistently reduced on the two datasets if the precision of contacts used with DeepRank is higher than 0.5, otherwise the impact of contacts is mixed.

### Section III. Successful cases of DeepRank in MULTICOM

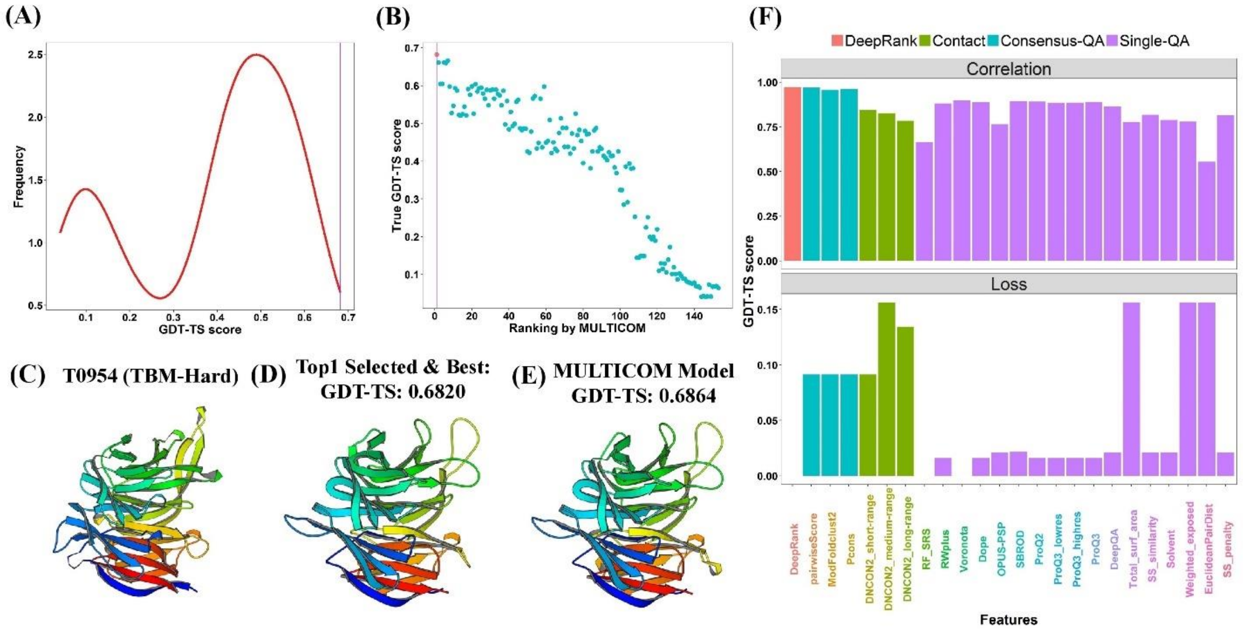


**Figure S7.** Tertiary structure prediction for T0954. **(A)** The distribution of GDT-TS scores of 153 server models. **(B)** The plot of the true GDT-TS scores of models against their predicted ranking by MULTICOM. The point highlighted in red is the top model selected by DeepRank. **(C)** The native structure of target T0954 (PDB code: 6cvz). **(D)** The top selected model. **(E)** The final first MULTICOM model (TS1). **(F)** The ranking of individual QA methods for target T0954.

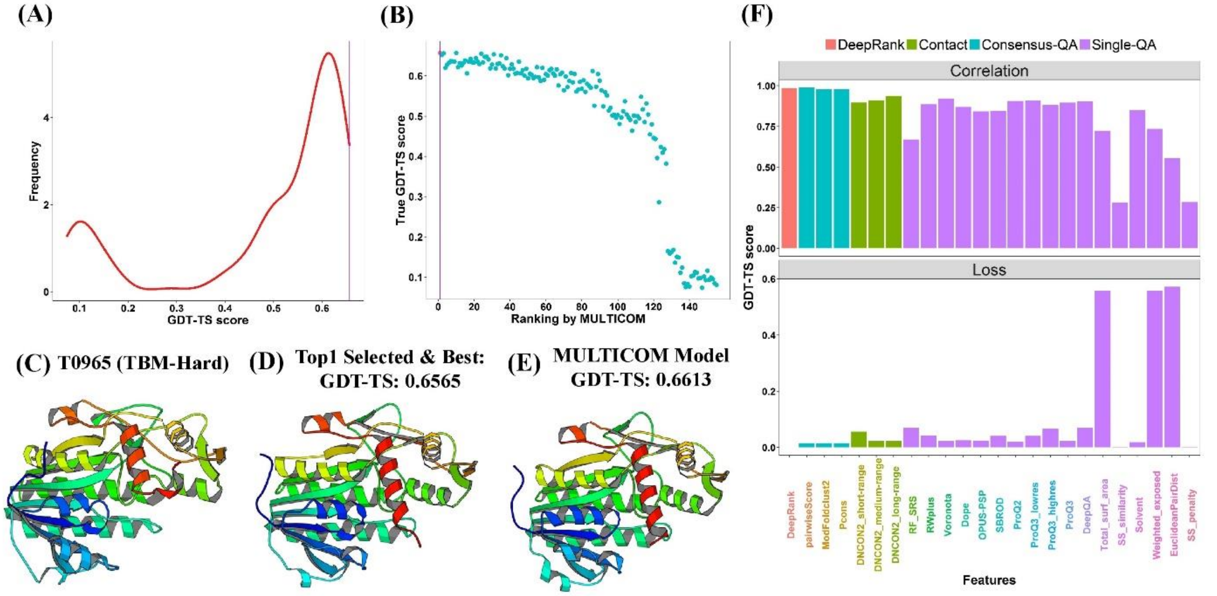


**Figure S8.** Tertiary structure prediction for T0965. **(A)** The distribution of GDT-TS scores of 155 server models. **(B)** The plot of the true GDT-TS scores of models against their predicted ranking by MULTICOM. The point highlighted in red is the top model selected by DeepRank. **(C)** The native structure of target T0965 (PDB code: 6d2v). **(D)** The top selected model. **(E)** The final first MULTICOM model (TS1)**. (F)** The ranking of individual QA methods for target T0965.

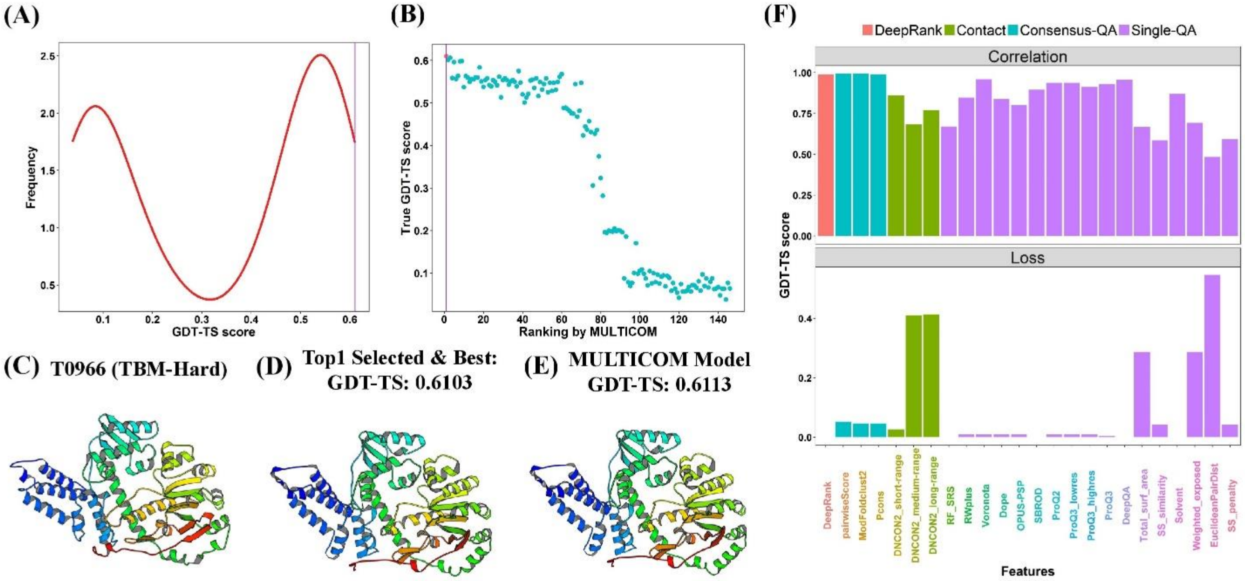


**Figure S9.** Tertiary structure prediction for T0966. **(A)** The distribution of GDT-TS scores of 146 server models. **(B)** The plot of the true GDT-TS scores of models against their predicted ranking by MULTICOM. The point highlighted in red is the top model selected by DeepRank. **(C)** The native structure of target T0966 (PDB code: 5w6l). **(D)** The top selected model. **(E)** The final first MULTICOM model (TS1). **(F)** The ranking of individual QA methods for target T0966.

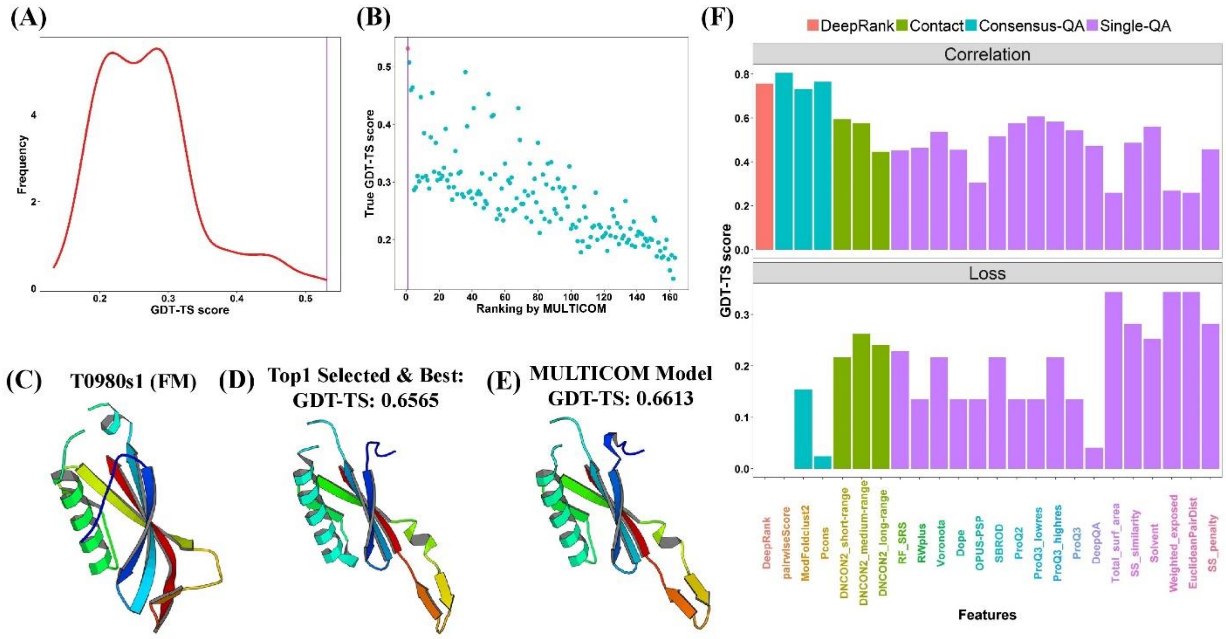

**Figure S10.** Tertiary structure prediction for T0980s1. **(A)** The distribution of GDT-TS scores of 163 server models. **(B)** The plot of the true GDT-TS scores of models against their predicted ranking by MULTICOM. The point highlighted in red is the top model selected by DeepRank. **(C)** The native structure of target T0980s1 (PDB code: 6gnx). **(D)** The top selected model. **(E)** The final first MULTICOM model (TS1). **(F)** The ranking of individual QA methods for target T0980s1.

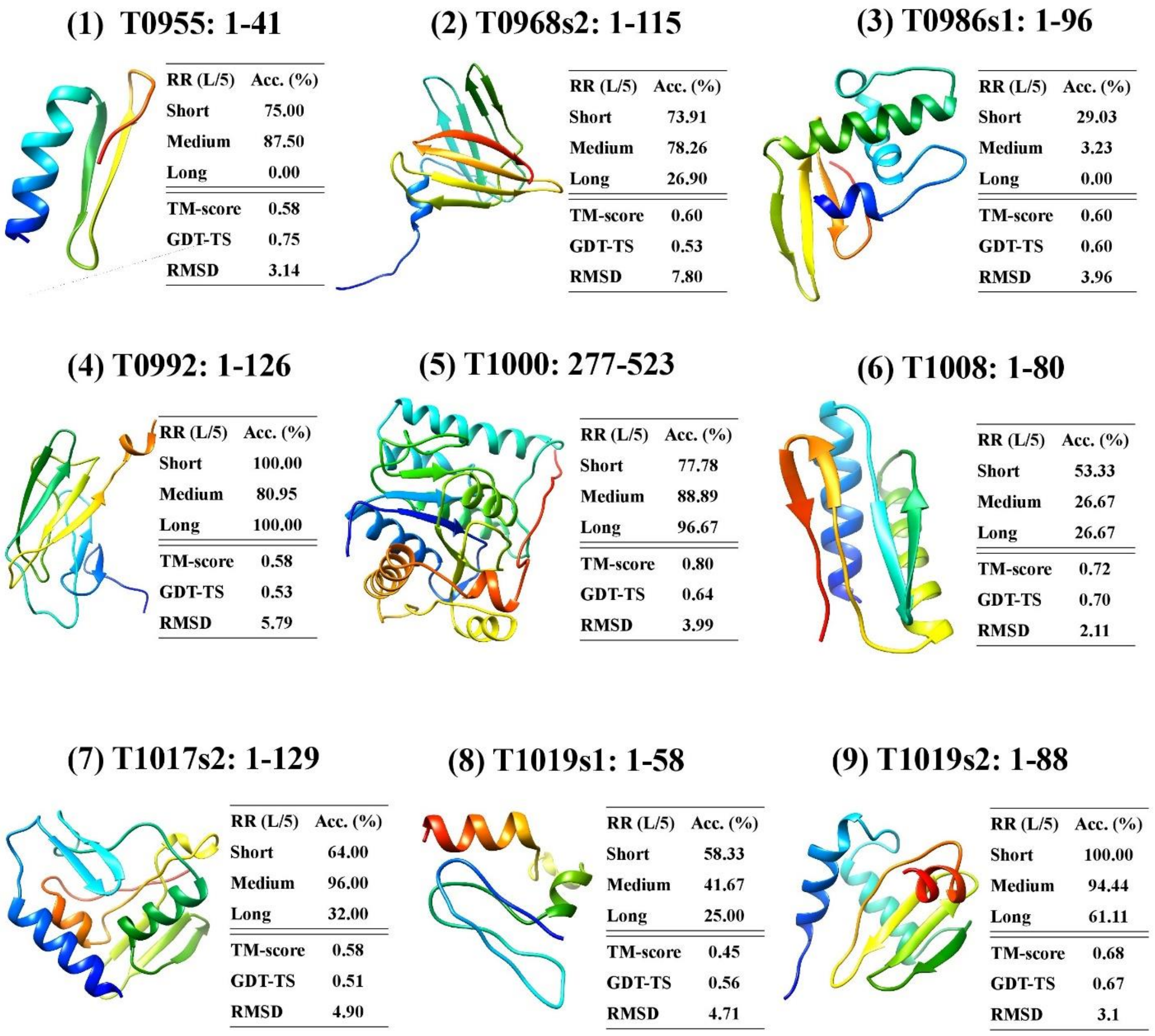


**Figure S11.** Examples of structural models built from predicted contacts for nine hard targets. The region that was modeled using contacts is visualized for each target. The accuracy of predicted top L/5 contacts of short-range, medium-range and long-range is reported in the table next to each model. The quality scores of each model are also provided. The experimental structures of these targets are not officially released and therefore not shown.

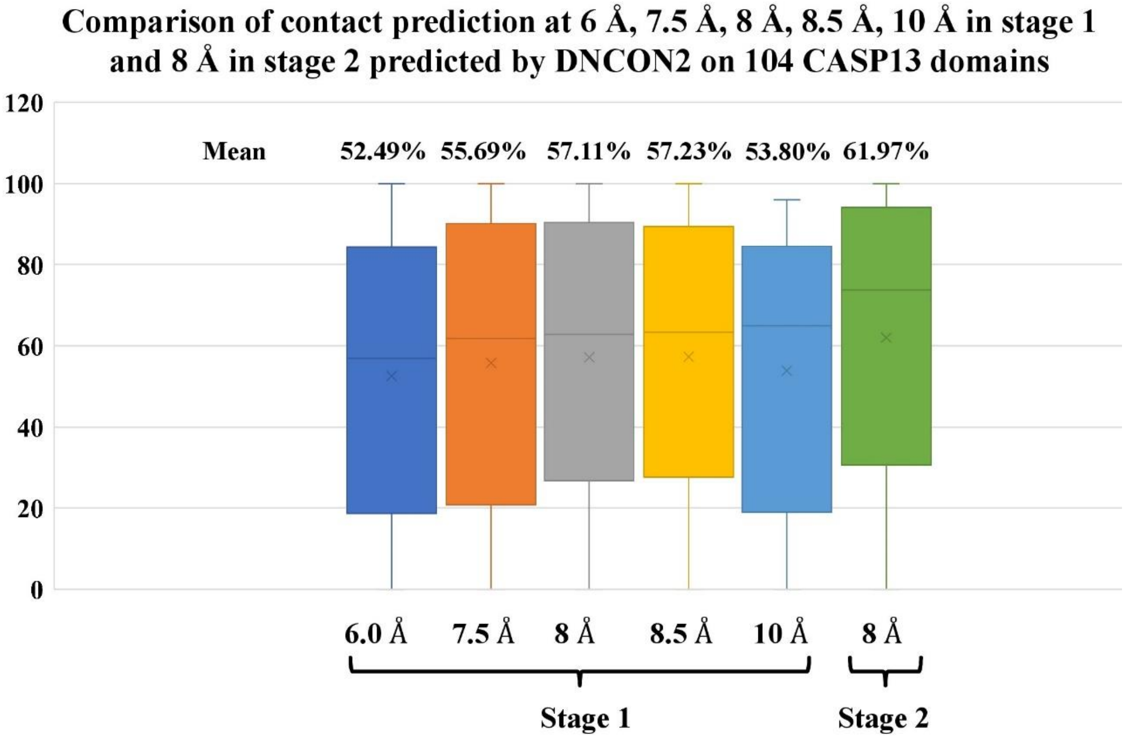


**Figure S12.** Comparison of contact prediction at 6 Å, 7.5 Å, 8 Å, 8.5 Å, 10 Å in stage 1 and 8 Å in stage 2 predicted by DNCON2. The accuracy of top L/5 predicted contact on two stages for 104 CASP13 "all groups" domains was analyzed and compared.

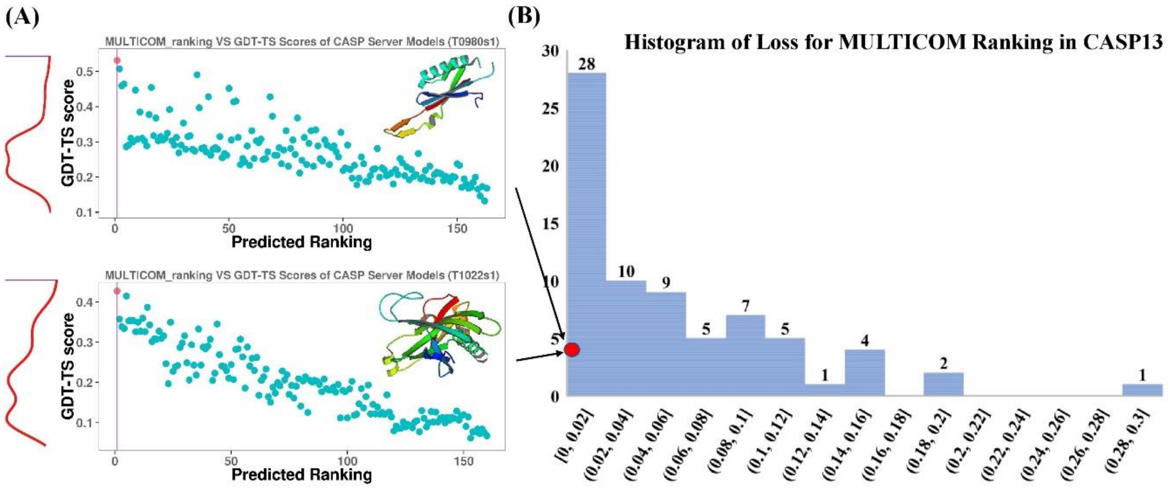


**Figure S13.** Performance of DeepRank method in CASP13. (A) Two good examples (T0980s1 and T1022s1) for which MULTICOM selected best server models (red dots) from the model pool.

(B) Histogram of loss for DeepRank method on CASP13 server models of all the targets. The bin size of loss is set to 0.02. The number of targets for each loss bin is show at the top.

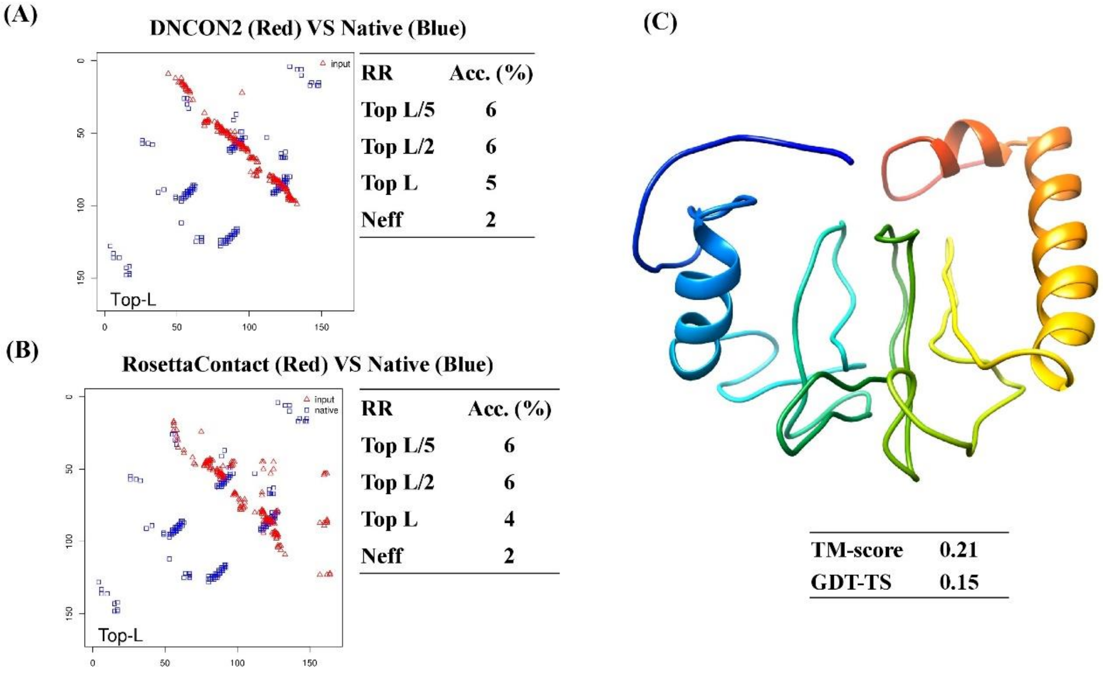


**Figure S14**. Failure of predicting and using contacts in modeling for target T0998. (A) Predicted contact map (red) versus true map (blue), accuracy of top L/5, L/2 and L predicted contacts, number of effective sequences (Neff =2). (B) The contact map of the model built by Rosetta with contacts (red) versus true contact map (blue) and accuracy of the contacts in the model. (C) The predicted structure using RosettaContact and the quality of the model. The contact prediction accuracy is low and the quality of the model is poor. The experimental structure of this target is not officially released and therefore not shown.

### Section IV. Failed cases for DeepRank in MULITCOM

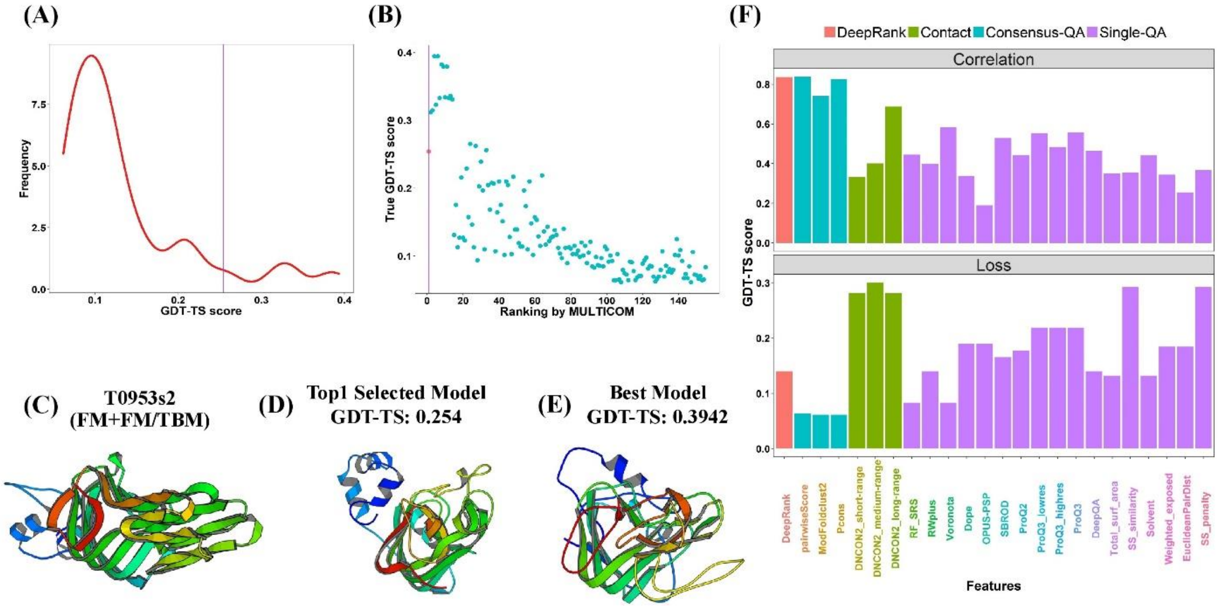


**Figure S15.** DeepRank model quality assessment for T0953s2. **(A)** The distribution of GDT-TS scores of 155 server models. **(B)** The plot of the true GDT-TS scores of models against their predicted ranking by MULTICOM. The point highlighted in red is the top model selected by DeepRank. **(C)** The native structure of target T0953s2 (PDB code: 6f45). **(D)** The top selected model. **(E)** The best server model. **(F)** The ranking of individual QA methods for target T0953s2.

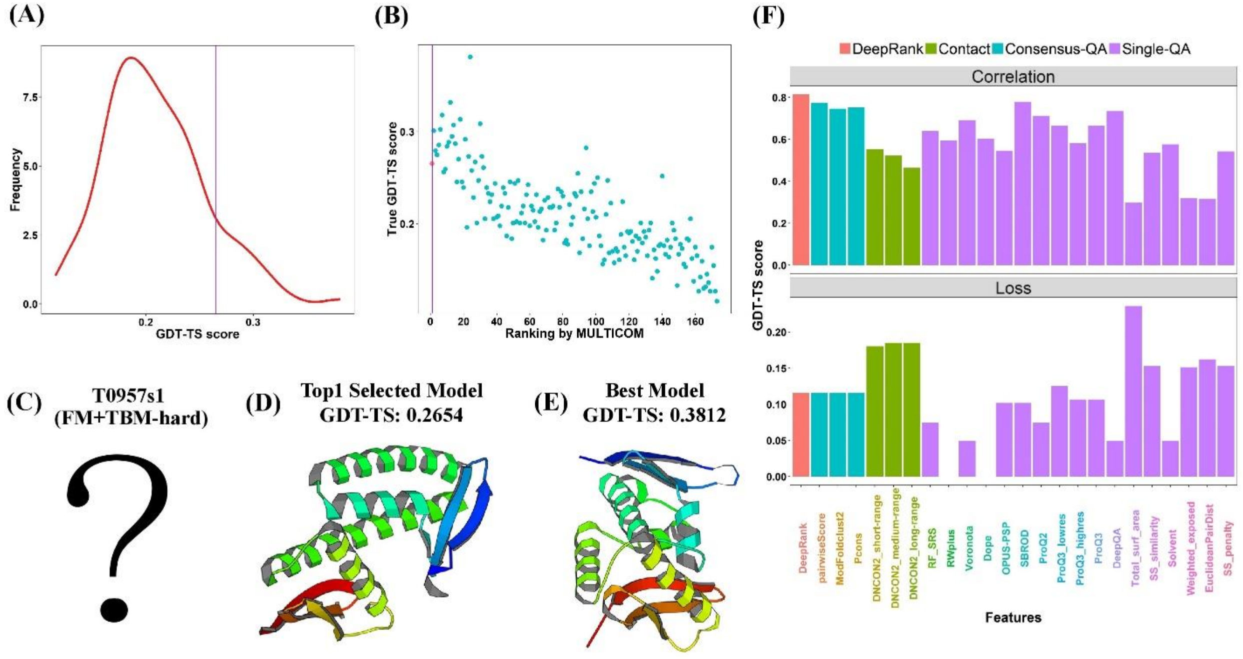

**Figure S16.** DeepRank model quality assessment for T0957s1. **(A)** The distribution of GDT-TS scores of 173 server models. **(B)** The plot of the true GDT-TS scores of models against their predicted ranking by MULTICOM. The point highlighted in red is the top model selected by DeepRank. **(C)** The experimental structure of target T0957s1 was not officially released to date. **(D)** The top selected model. **(E)** The best server model. **(F)** The ranking of individual QA methods for target T0957s1.

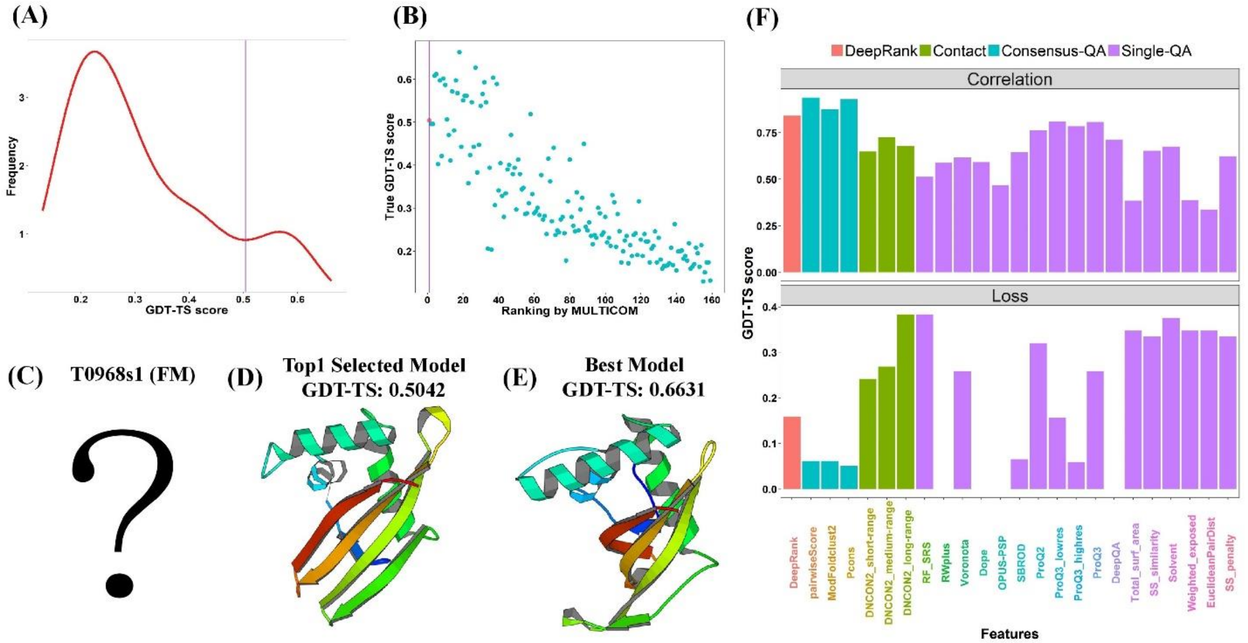


**Figure S17.** DeepRank model quality assessment for T0968s1. **(A)** The distribution of GDT-TS scores of 159 server models. **(B)** The plot of the true GDT-TS scores of models against their predicted ranking by MULTICOM. The point highlighted in red is the top model selected by DeepRank. **(C)** The experimental structure of target T0968s1 was not officially released to date. **(D)** The top selected model. **(E)** The best server model. **(F)** The ranking of individual QA methods for target T0968s1.

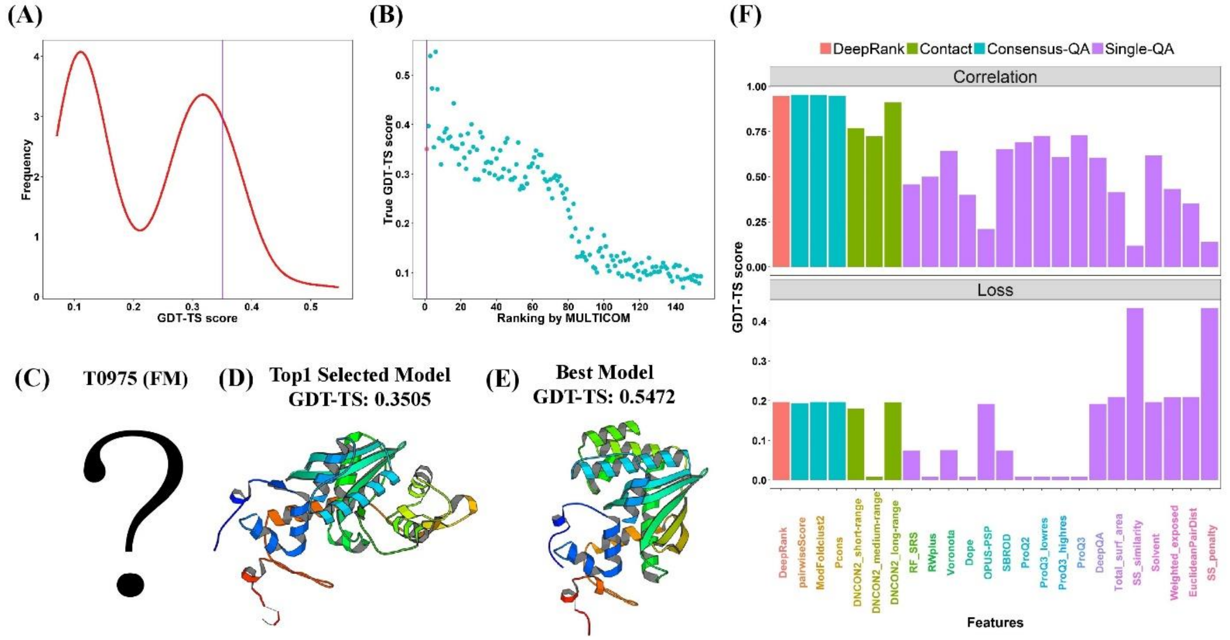


**Figure S18.** DeepRank model quality assessment for T0975. **(A)** The distribution of GDT-TS scores of 154 server models. **(B)** The plot of the true GDT-TS scores of models against their predicted ranking by MULTICOM. The point highlighted in red is the top model selected by DeepRank. **(C)** The experimental structure of target T0975 was not officially released to date. **(D)** The top selected model. **(E)** The best server model. **(F)** The ranking of individual QA methods for target T0975.

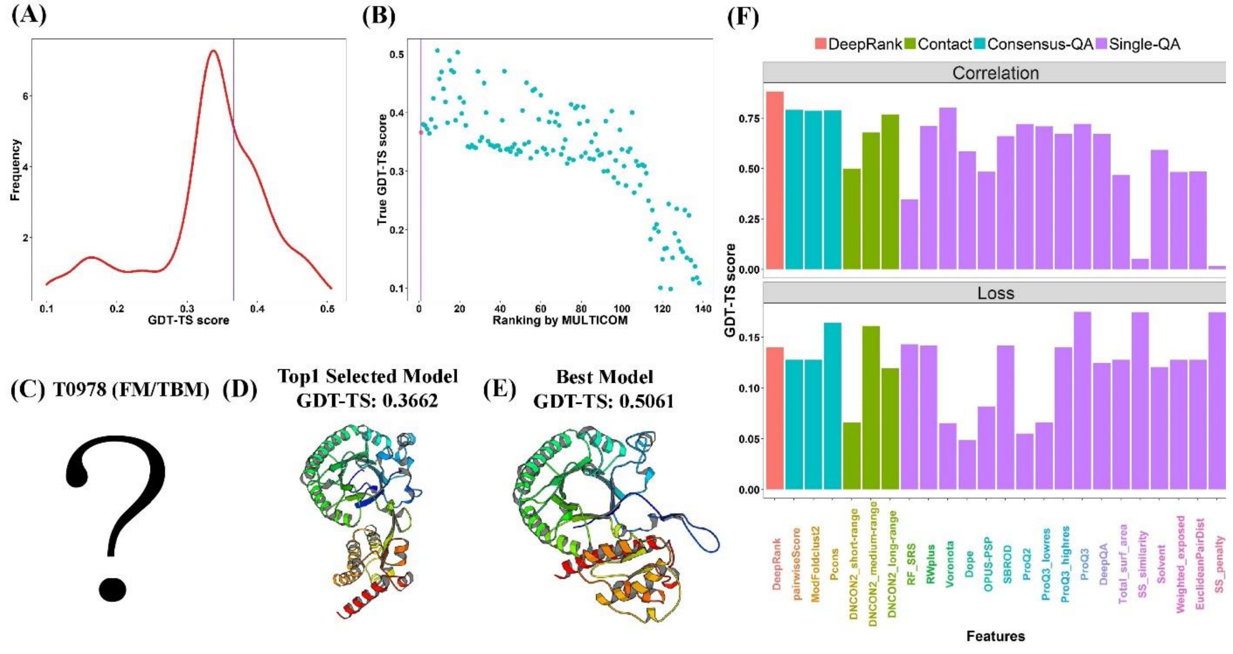


**Figure S19.** DeepRank model quality assessment for T0978. **(A)** The distribution of GDT-TS scores of 138 server models. **(B)** The plot of the true GDT-TS scores of models against their predicted ranking by MULTICOM. The point highlighted in red is the top model selected by DeepRank. **(C)** The experimental structure of target T0978 was not officially released to date. **(D)** The top selected model. **(E)** The best server model. **(F)** The ranking of individual QA methods for target T0978.

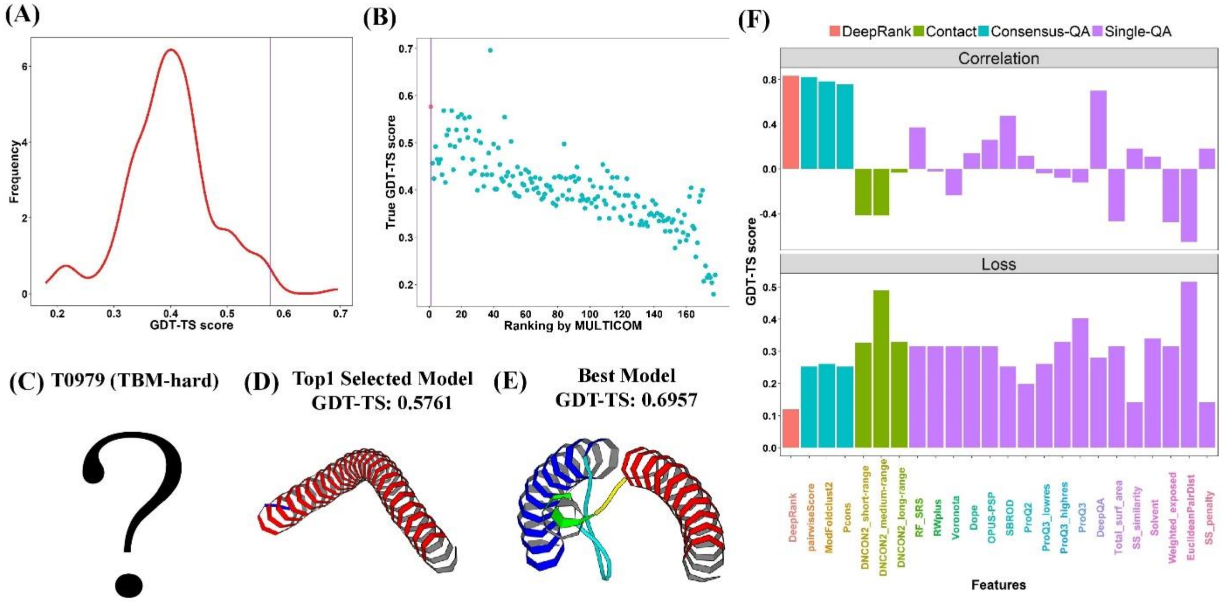


**Figure S20.** DeepRank model quality assessment for T0979. **(A)** The distribution of GDT-TS scores of 178 server models. **(B)** The plot of the true GDT-TS scores of models against their predicted ranking by MULTICOM. The point highlighted in red is the top model selected by DeepRank. **(C)** The experimental structure of target T0979 was not officially released to date. **(D)** The top selected model. **(E)** The best server model. **(F)** The ranking of individual QA methods for target T0979.

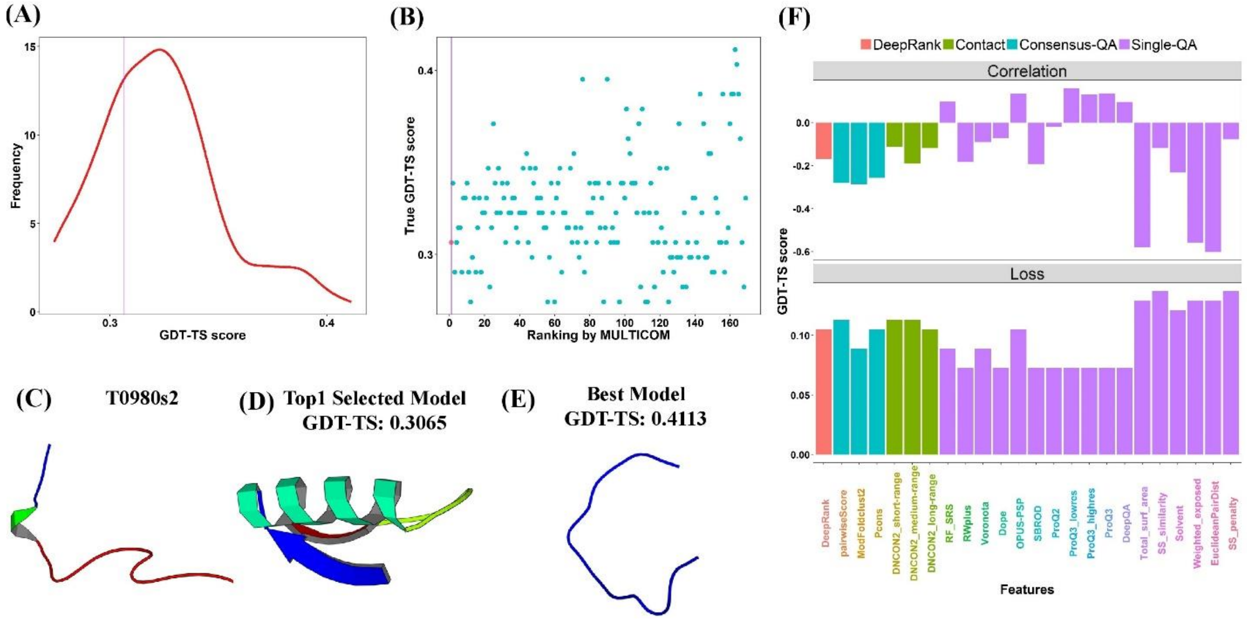

**Figure S21.** DeepRank model quality assessment for T0980s2. **(A)** The distribution of GDT-TS scores of 169 server models. **(B)** The plot of the true GDT-TS scores of models against their predicted ranking by MULTICOM. The point highlighted in red is the top model selected by DeepRank. **(C)** The native structure of target T0980s2 (PDB code: 6qnx). **(D)** The top selected model. **(E)** The best server model. **(F)** The ranking of individual QA methods for target T0980s2.

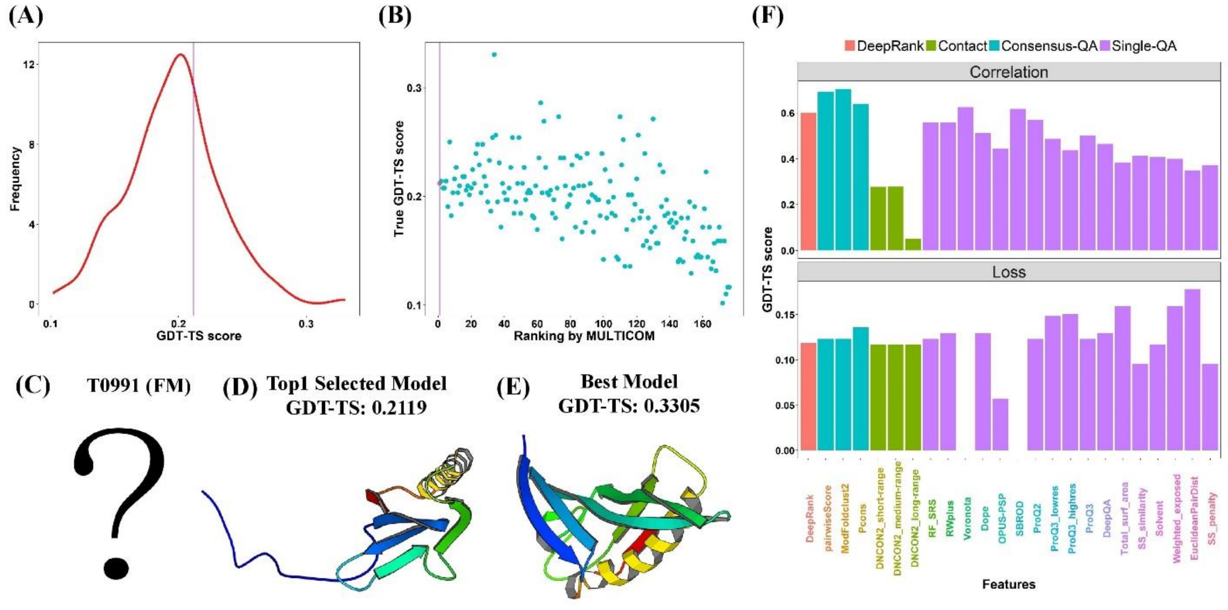

**Figure S22.** DeepRank model quality assessment for T0991. **(A)** The distribution of GDT-TS scores of 176 server models. **(B)** The plot of the true GDT-TS scores of models against their predicted ranking by MULTICOM. The point highlighted in red is the top model selected by DeepRank. **(C)** The experimental structure of target T0991 was not officially released to date. **(D)**

The top selected model. **(E)** The best server model. **(F)** The ranking of individual QA methods for target T0991.

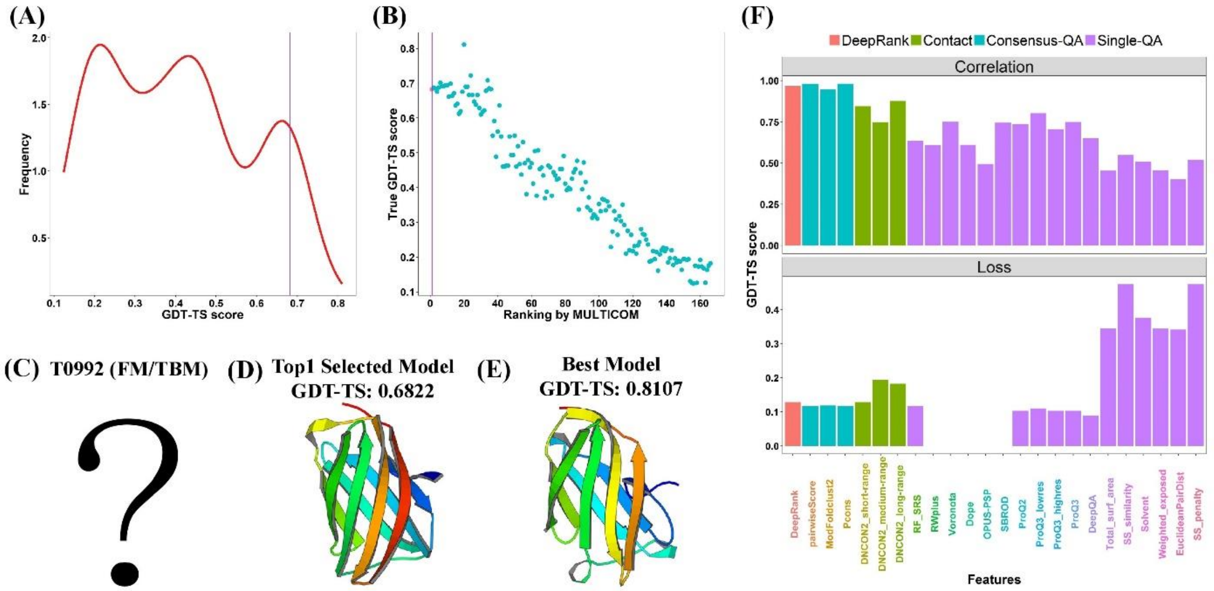


**Figure S23.** DeepRank model quality assessment for T0992. **(A)** The distribution of GDT-TS scores of 166 server models. **(B)** The plot of the true GDT-TS scores of models against their predicted ranking by MULTICOM. The point highlighted in red is the top model selected by DeepRank. **(C)** The experimental structure of target T0992 was not officially released to date. **(D)** The top selected model. **(E)** The best server model. **(F)** The ranking of individual QA methods for target T0992.

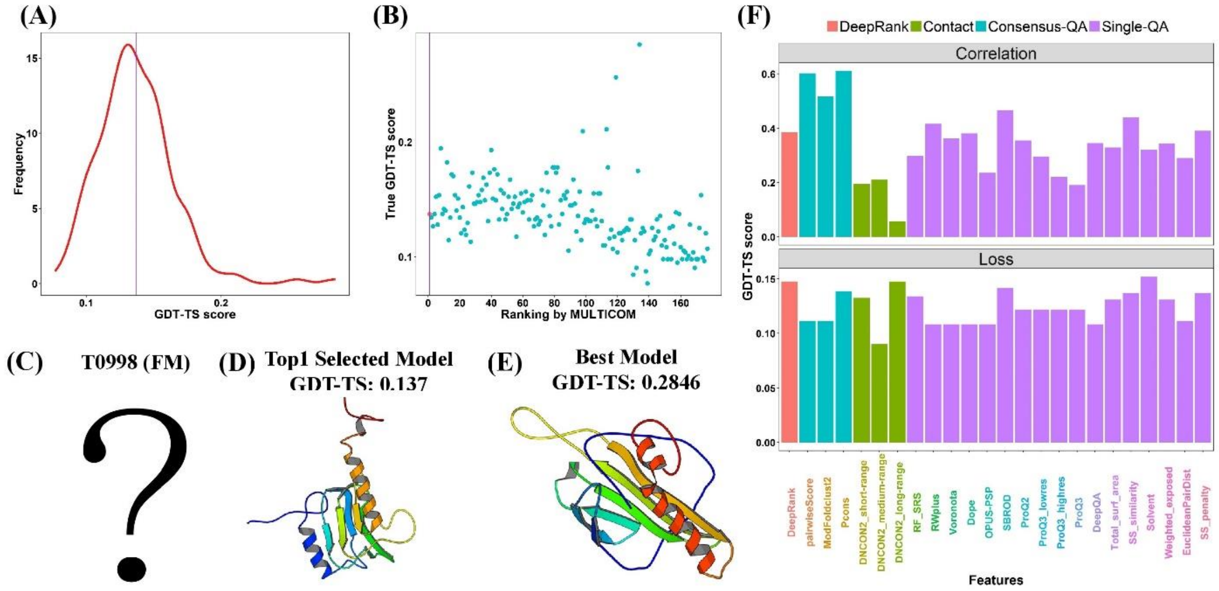

**Figure S24.** DeepRank model quality assessment for T0998. **(A)** The distribution of GDT-TS scores of 177 server models. **(B)** The plot of the true GDT-TS scores of models against their predicted ranking by MULTICOM. The point highlighted in red is the top model selected by DeepRank. **(C)** The experimental structure of target T0998 was not officially released to date. **(D)** The top selected model. **(E)** The best server model. **(F)** The ranking of individual QA methods for target T0998.

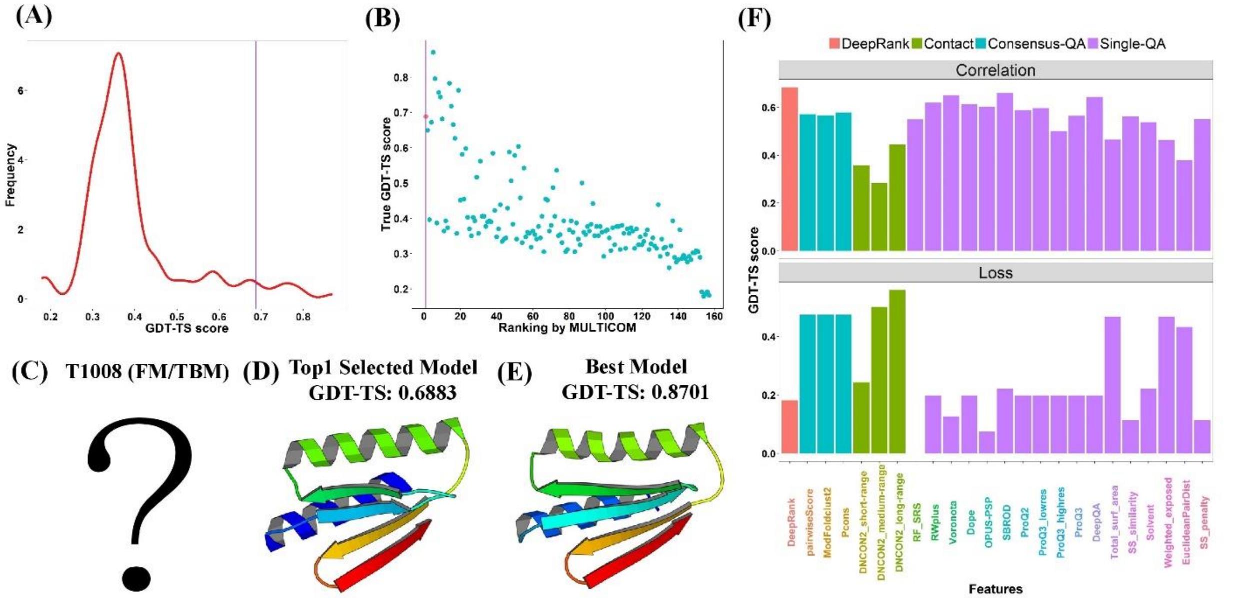


**Figure S25.** DeepRank model quality assessment for T1008. **(A)** The distribution of GDT-TS scores of 157 server models. **(B)** The plot of the true GDT-TS scores of models against their predicted ranking by MULTICOM. The point highlighted in red is the top model selected by DeepRank. **(C)** The experimental structure of target T1008 was not officially released to date. **(D)** The top selected model. **(E)** The best server model. **(F)** The ranking of individual QA methods for target T1008.

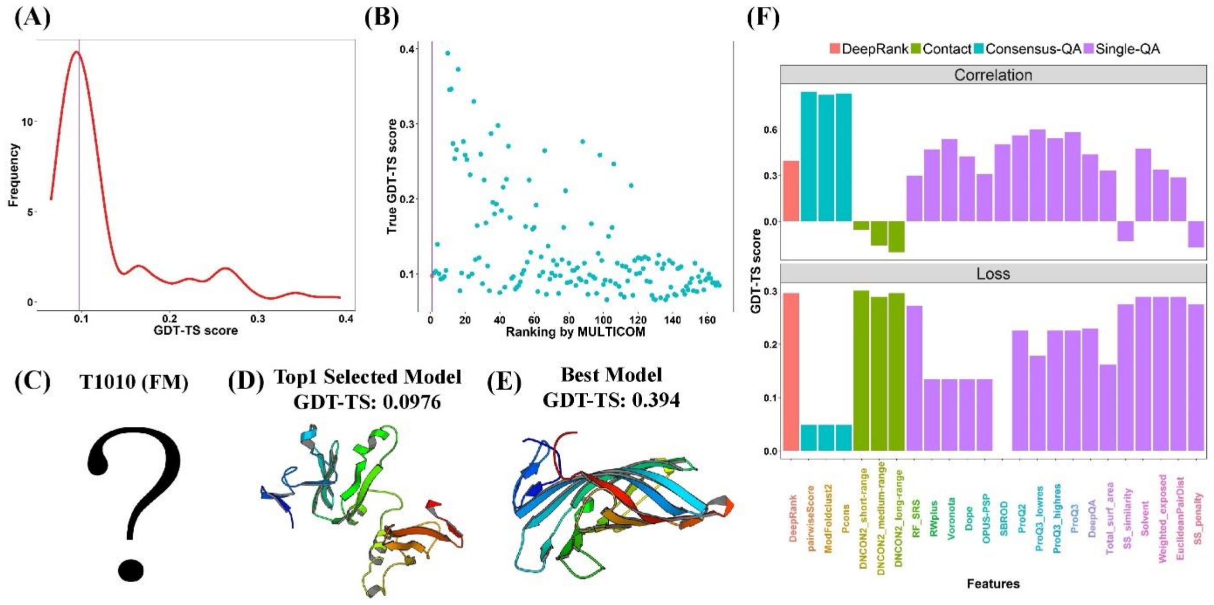


**Figure S26.** DeepRank model quality assessment for T1010. **(A)** The distribution of GDT-TS scores of 167 server models. **(B)** The plot of the true GDT-TS scores of models against their predicted ranking by MULTICOM. The point highlighted in red is the top model selected by DeepRank. **(C)** The experimental structure of target T1010 was not officially released to date. **(D)** The top selected model. **(E)** The best server model. **(F)** The ranking of individual QA methods for target T1010.

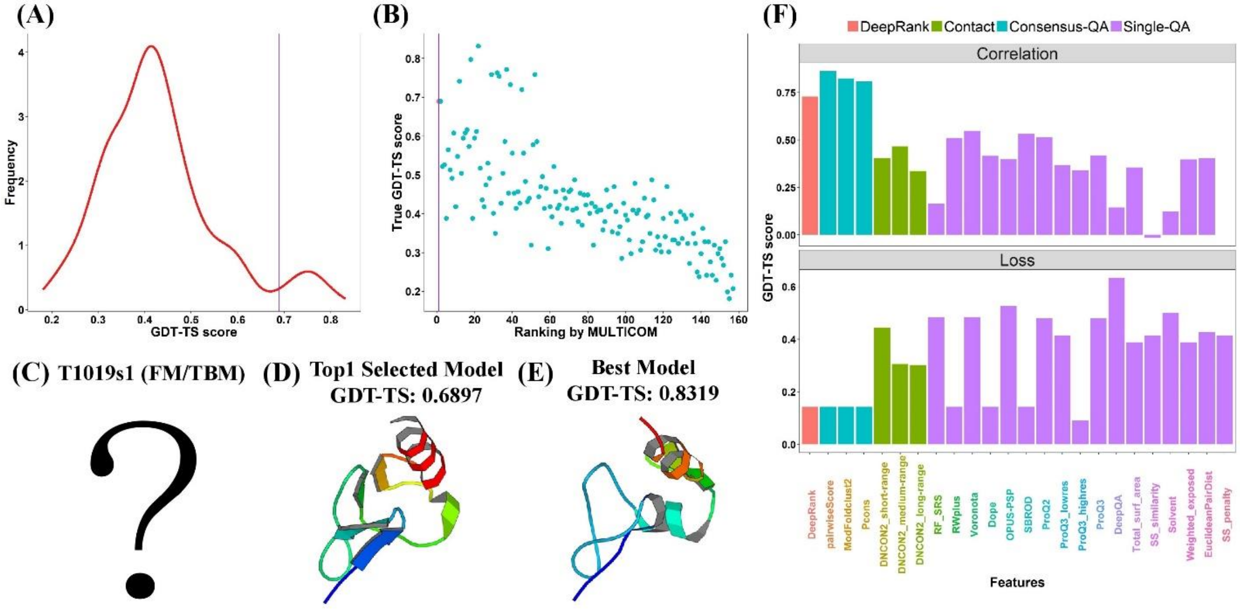


**Figure S27.** DeepRank model quality assessment for T1019s1. **(A)** The distribution of GDT-TS scores of 157 server models. **(B)** The plot of the true GDT-TS scores of models against their predicted ranking by MULTICOM. The point highlighted in red is the top model selected by DeepRank. **(C)** The experimental structure of target T1019s1 was not officially released to date.

**(D)** The top selected model. **(E)** The best server model. **(F)** The ranking of individual QA methods for target T1019s1.

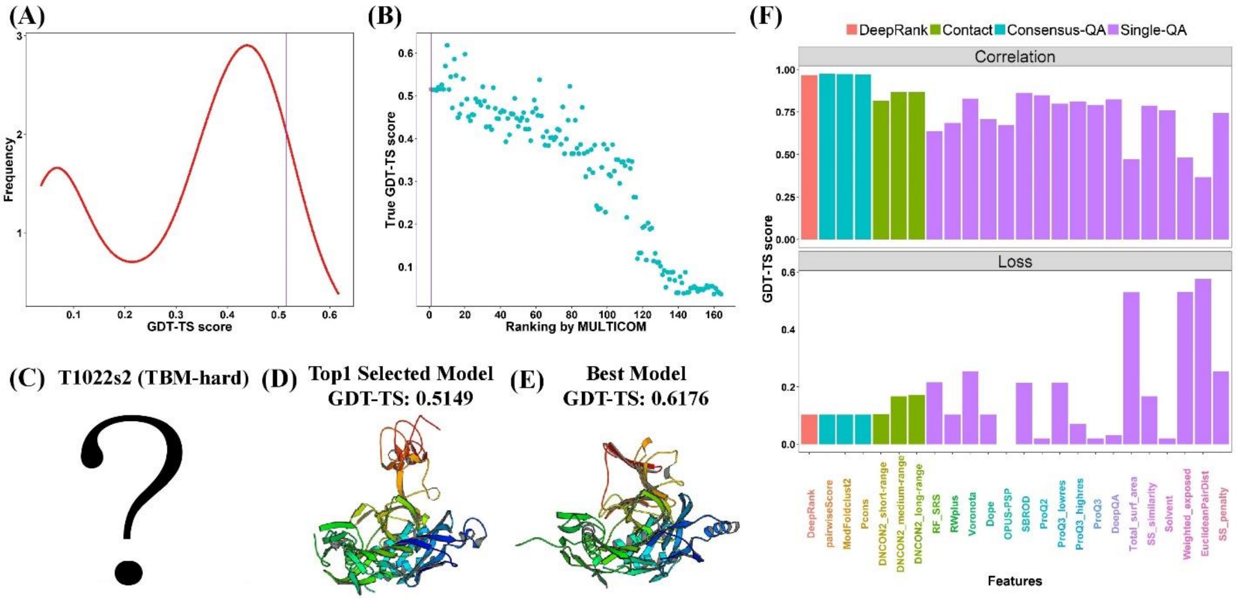


**Figure S28.** DeepRank model quality assessment for T1022s2. **(A)** The distribution of GDT-TS scores of 164 server models. **(B)** The plot of the true GDT-TS scores of models against their predicted ranking by MULTICOM. The point highlighted in red is the top model selected by DeepRank. **(C)** The experimental structure of target T1022s2 was not officially released to date. **(D)** The top selected model. **(E)** The best server model. **(F)** The ranking of individual QA methods for target T1022s2.
